# CK2α phosphorylates CHIP for oncogenesis: A novel post-translational switch in cancer

**DOI:** 10.1101/2025.10.15.682604

**Authors:** Subhajit Karmakar, Mouli Chatterjee, Malini Basu, Mrinal K Ghosh

**Affiliations:** Cancer Biology and Inflammatory Disorder Division, Council of Scientific and Industrial Research-Indian Institute of Chemical Biology (CSIR-IICB), TRUE Campus, CN-6, Sector– V, Salt Lake, Kolkata-700091 & 4, Raja S.C. Mullick Road, Jadavpur, Kolkata-700032, India; Academy of Scientific and Innovative Research (AcSIR), Ghaziabad, Uttar Pradesh-201002, India; Department of Microbiology, Dhruba Chand Halder College, University of Calcutta, Dakshin Barasat, WB, India

**Keywords:** CK2α, CHIP, Phosphorylation, Degradation, Post-translational modification (PTM), Cancer

## Abstract

The post-translational regulation of tumor suppressors by oncogenic kinases remains a critical yet underexplored determinant of proteostasis reprogramming in cancer. Here, we identified the E3 ubiquitin ligase CHIP (C-terminus of Hsc70-Interacting Protein) as a direct and functionally relevant substrate of the serine/threonine kinase CK2α, which is frequently overexpressed in solid tumors. Using LC/MS analysis, *in silico* kinase prediction, and molecular interaction mapping, we demonstrated that CK2α phosphorylates CHIP at serine 19 (represented as S19), a conserved residue within its TPR domain, thereby promoting ubiquitination and subsequent proteasomal degradation of CHIP. This phosphorylation-dependent destabilization of CHIP impairs its ability to target oncogenic substrates, such as AKT, resulting in sustained AKT phosphorylation and activation, required for oncogenesis. Clinically, we observed a robust inverse correlation between CK2α and CHIP expressions across colorectal and breast cancer patient datasets, which was validated by immunofluorescence (IF) analyses in tumor samples. Multiple functional assays revealed that CK2α suppression, either by genetic ablation or pharmacological inhibition by TBCA, restores CHIP stability, reactivates apoptotic signalling cascades, and attenuates tumor cell proliferation, migration, and 3D spheroid integrity. Moreover, CK2α blockade in syngeneic murine models diminishes primary tumor burden and metastatic dissemination, concomitant with increased CHIP accumulation and reduced AKT signalling. Mechanistically, a phosphorylation-resistant mutant of CHIP at S19 (CHIP-S19A), is refractory to CK2α- mediated degradation and preserves anti-tumor functions, delineating a phosphorylation-dependent proteolytic switch as a central node in the ‘CK2α-CHIP-AKT’ regulatory axis. These findings establish a novel paradigm wherein an oncogenic kinase hijacks protein quality control pathway to suppress tumor suppressive activity and facilitate malignancy. Our study positions CK2α as a druggable modulator of CHIP turnover, offering a translational framework for restoring proteostatic checkpoints and constraining oncogenic signalling in aggressive cancers.

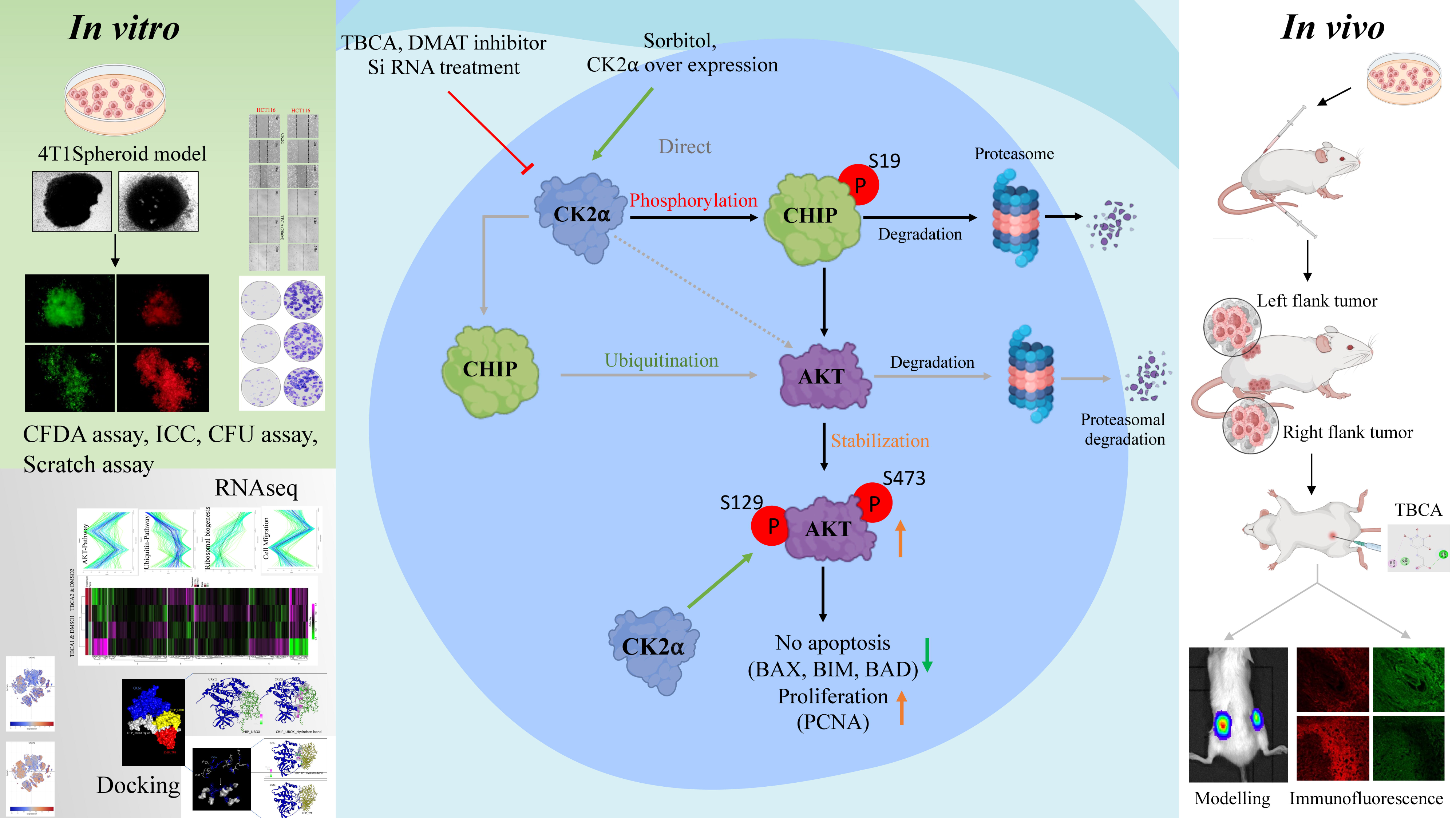

## INTRODUCTION

The maintenance of cellular homeostasis relies on the coordinated regulation of signal transduction pathways and proteostatic mechanisms, which collectively govern protein stability, function, and turnover. Within this regulatory framework, the E3 ubiquitin ligase CHIP (encoded by *STUB1*), a co-chaperone, has emerged as a pivotal node linking the molecular chaperone machinery to the ubiquitin-proteasome system (UPS) [1]. By facilitating the polyubiquitination and degradation of misfolded or damaged proteins. CHIP enforces a quality-control checkpoint that is critical not only for normal cellular physiology but also for the attenuation of oncogenic signalling. Despite its reported tumor-suppressive function in diverse malignancies [2–8], the upstream mechanisms that modulate CHIP expression, its post-translational regulation and turnover remain incompletely understood. Casein kinase 2 (CK2), a constitutively active serine/threonine kinase, is frequently upregulated in a wide spectrum of malignancies and has emerged as a central player in modulating cellular processes, such as proliferation, apoptosis resistance, and metabolic rewiring [9, 10]. Although CK2α, the catalytic subunit of the CK2 tetramer has been implicated in the phosphorylation of numerous oncogenic substrates [11–14], its role in regulating CHIP, a tumor suppressor that controls AKT stability and activity, has remained unexplored [15–17].

In this study, we identified CHIP as a novel substrate of CK2α. We demonstrated that CK2α phosphorylates CHIP at S19, thereby marking it for ubiquitination and subsequent proteasomal degradation. This phosphorylation-dependent mechanism functionally silences tumor-suppressive capacity of CHIP, stabilizes oncogenic AKT signalling, and promotes tumor growth and metastasis. Through a combination of mass spectrometry, mutational analysis, proteogenomic correlation, 3D spheroid models, and *in vivo* tumor xenografts, we define the ‘CK2α-CHIP-AKT’ axis as a mechanistically and clinically relevant pathway in breast and colorectal cancers. Pharmacological inhibition of CK2α using TBCA rescues CHIP expression, restores apoptosis, and constrains tumor progression.

Collectively, our findings uncover a post-translational mechanism by which CK2α suppresses CHIP’s function to facilitate oncogenic signalling, revealing a targetable vulnerability in cancers. Therefore, we are showing for the first time, CK2α dependent phosphorylation of CHIP at the S19 residue primes it for proteasomal degradation, leading to AKT stabilization and cancer progression.

## RESULTS

### CK2α colocalizes and interacts with CHIP and phosphorylates at S19: Identification of a novel phosphorylation site

Previous proteomic studies identified multiple CHIP phosphorylation sites but lacked details on responsible kinases, phosphorylation-dependent degradation pathways, or physiological responses [18, 19]. To define CHIP regulation, we investigated whether CK2α, a serine/threonine kinase overexpressed in cancer, phosphorylates CHIP and modulates its stability. Bioinformatics analysis (KinasePhos, NetPhos 3.1, ScanSite) identified S19 (S20 in mouse) as a CK2α consensus site (S/T-X-X-D/E) previously reported to stabilize CHIP activity. Our findings suggest CK2α-mediated S19 phosphorylation promotes CHIP degradation, representing a novel mechanism influencing stability and protein homeostasis. Putative CK2α sites were located within the conserved C-terminal TPR domain of CHIP, corresponding to an unstructured solvent-exposed region, as expected for CHIP sequences (Fig. 1A, Supplementary Fig. 1A, B). To examine CK2α-dependent regulation of CHIP, we analyzed their interaction by immunoprecipitation (IP) and silver staining in HEK293 cells (Fig. 1B). Given the transient dynamics of the enzymatic reaction, the interaction between overexpressed CHIP and CK2α was further monitored by forward & reverse IP experiments (Fig. 1C). To validate the CK2α-CHIP interaction, their colocalization was also investigated by using immunofluorescence (IF) in HEK293, HCT116, MDA-MB-231, MDA-MB-468, and MCF-7 cells. Both the proteins showed prominent cytoplasmic and nuclear colocalization with high Pearson’s correlation coefficients (R=0.90, 0.75, 0.85, 0.81, and 0.94, respectively). These results indicate a robust association between CK2α and CHIP, suggesting possible formation of a functional complex (Fig. 1D).

**Fig. 1.**
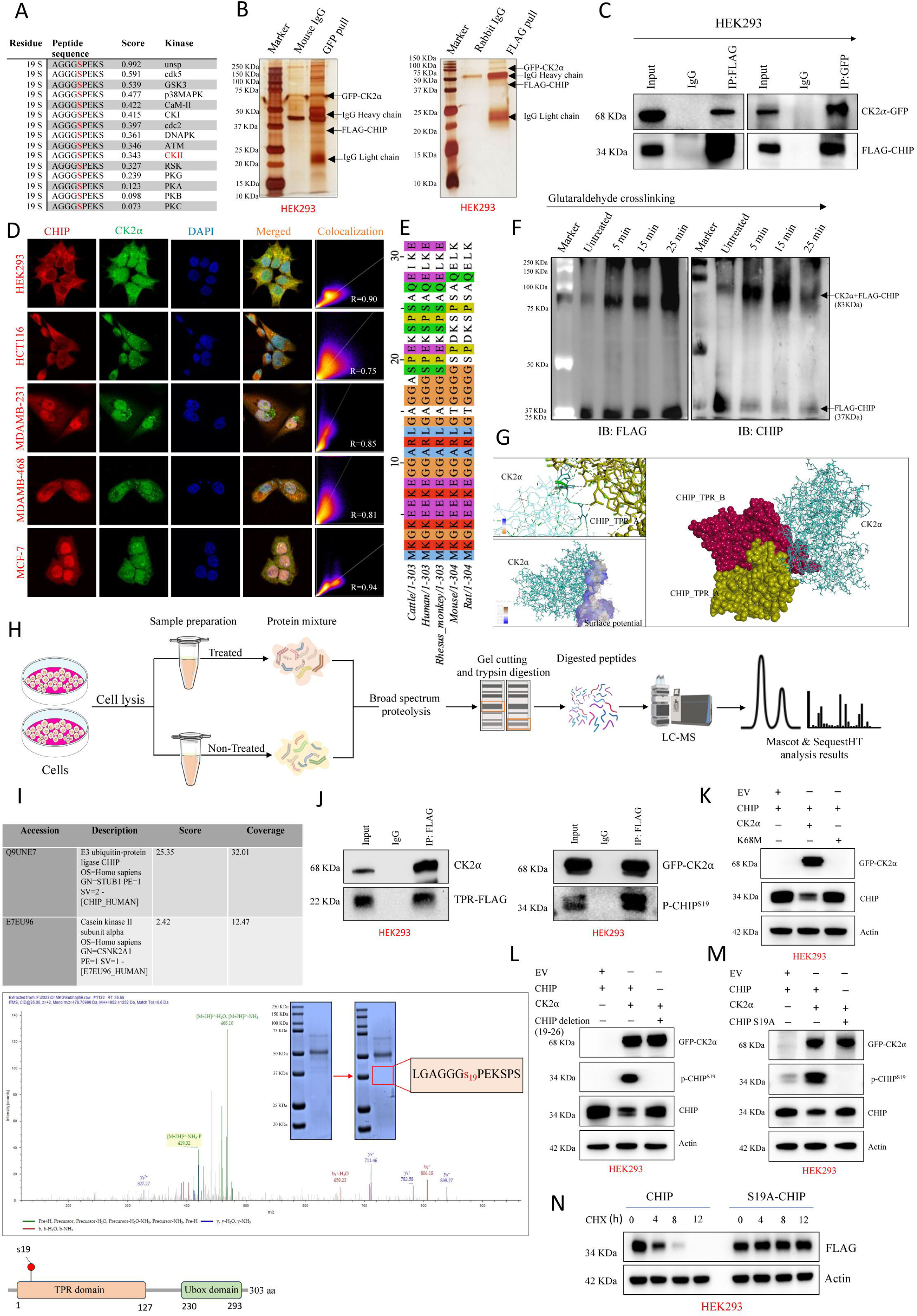
CK2α phosphorylates CHIP at serine 19 to regulate its stability through a phosphorylation-dependent degradation mechanism. **A** Putative CK2α phosphorylation sites within CHIP, located near the conserved TPR domain. **B-C** Immunoprecipitation (IP) and silver staining showing interaction between CK2α and CHIP in HEK293 cells. **D** Immunofluorescence (IF) in multiple cell lines (HEK293, HCT116, MDA-MB-231, MDA-MB-468, MCF-7) demonstrating cytoplasmic and nuclear colocalization of CK2α and CHIP. **E** Multiple sequence alignment of CHIP’s TPR domains across species highlighting conserved residues as potential CK2α-binding sites. **F** Glutaraldehyde cross-linking and immunoblotting (IB) confirming CHIP-CK2α complex formation. **G** Molecular docking analysis of CHIP-CK2α interaction interfaces, visualized in PyMOL. **H-I** LC-MS proteomic analysis identifying CHIP and CK2α with significant coverage and detection of CHIP’s phosphopeptide. **J** Domain mapping confirming TPR domain of CHIP as the primary CK2α interaction site. **K** Co-expression studies with wild-type and mutant CK2α-K68M validating phosphorylation dependency. **L-M** IB with CHIP mutants (S19A, Δ19-26) and phospho-specific antibodies confirming CK2α-dependent phosphorylation at serine 19. **N** Figure showing CHIP-S19A mutant is more stable than its wild-type form.

Next, multiple sequence alignment of the tetratricopeptide repeat (TPR) domain of CHIP from cattle, humans, monkeys, mice, and rats was performed using Jalview software (Fig. 1E). Sequence alignment revealed some conserved amino acid residues in CHIP, identified as potential CK2α-phosphorylation sites, and the regions were important for their interaction. To further confirm the CHIP-CK2α interaction, glutaraldehyde cross-linking and immunoblotting (IB) was performed with whole cell lysates (WCL) prepared from cells overexpressing both FLAG-CHIP and GFP-CK2α. Time-dependent cross-linking (0-25 min) produced a higher molecular weight band of the CHIP-CK2α complex, and immunoblotting (IB) with anti-FLAG antibody confirmed the presence of CHIP in this complex (Fig. 1F). To define the interfaces in CHIP-CK2α interaction molecular docking was conducted using the crystal structures of CK2α (PDB ID: 3AT2) and CHIP (PDB ID: 2C2L) from the RCSB Protein Data Bank. After structural refinement, the ClusPro docking server predicted binding residues and surface potentials, and the complex was visualized in PyMOL. This model provides structural insights into the CHIP-CK2α interaction, corroborating the biochemical data (Fig. 1G, Supplementary Fig. 1C).

The experimental workflow for proteomic analysis involves culturing and treating cells under defined conditions, followed by protein extraction, gel electrophoresis, and LC-MS. Data processed using Mascot and SequestHT enabled identification of protein-protein interactions, post-translational modifications (PTM), phosphorylation sites, and mapping of the CK2α-CHIP interaction network (Fig. 1H). LC-MS analysis identified two key proteins, CHIP (Q9UNE7) and CK2α (E7EU96), with association scores of 25.35 and 2.42, respectively. The coverage rates for CHIP and CK2α were 32.01% and 12.47%, respectively, indicating a significant presence of these proteins in the analyzed samples. In addition, the phosphopeptide sequence of CHIP was detected, suggesting that phosphorylation may play an important role in regulating CHIP and its functions (Fig. 1I). To validate the interaction between CK2α and CHIP, IP assays were performed. The results confirmed that CK2α associates with CHIP in a pull-down experiment, demonstrating their molecular interaction. Domain mapping of CHIP revealed that the TPR domain is the primary region responsible for its interaction with CK2α. This structural insight is crucial for CK2α-CHIP interaction and its functional implications in cancer cells (Fig. 1J). To further demonstrate this interaction, FLAG-tagged CHIP was co-expressed with either GFP-CK2α or HA-tagged CK2α-K68M, a previously described dominant-negative mutant unable to bind with ATP, hence no phosphorylation observed (Fig. 1K). Additionally, we supported our findings by IB using CHIP mutants *viz*, phospho-defective S19A point mutation (CHIP-S19A) and deletion of 19-26 residues, created by site-directed mutagenesis (SDM) followed by IB with P-S19CHIP antibody or PanS/T antibody (Fig. 1L-M, Supplementary Fig. 1D). These results suggest that CK2α regulates CHIP phosphorylation at a specific serine residue. Next, we wanted to check whether this phosphorylation by CK2α influences CHIP’s stability or not, and we observed that CHIP-S19A was more stable than its wild type form (Fig. 1N). Thus, we identified a novel phosphorylation dependent degradation mechanism of CHIP and our results showed that CHIP is a CK2α substrate. These insights provide a foundation for future studies aimed at targeting the CK2α-CHIP interaction as a potential therapeutic strategy for cancer treatment. Actin was kept as loading control in all the IB analysis.

### CK2α and CHIP maintain an inverse correlation in cancer: A clinically relevant regulatory axis

To explore the clinical significance of CK2α and CHIP in cancer progression, we investigated their expressions across tumor datasets and patient samples. The Cancer Proteogenomic Data Analysis Site (cProSite) is used as a platform for the visualization of proteomic datasets across cancers [20]. To examine the clinical relevance of CK2⍺ and CHIP in cancers, we examined their mRNAs (CSNK2A1 & STUB1) and proteins (CK2⍺ & CHIP) expression levels in patient samples (Fig. 2A, Supplementary Fig. 2A). Our OncoDB and UALCAN database analysis shows a negative correlation between CK2⍺ and CHIP in both CPTAC datasets; in Colon Adenocarcinoma (COAD) and Breast Cancer (BRCA) the correlation coefficient was -0.54 and -0.24 respectively, indicating their inverse relation (Fig. 2B, Supplementary Fig. 2B) [21, 22]. Our analysis revealed a robust and statistically significant negative correlation between them, CK2⍺ (*CSNK2A1*) and CHIP (*STUB1*), suggesting that this inverse association may play an important role in modulating cancer progression. In contrast, no significant correlation was observed at the mRNA level, implying that CK2⍺-mediated regulation of CHIP is likely governed by post-translational mechanisms in breast and colon cancers (Supplementary Fig. 2C). Next, we performed single-cell transcriptomic analysis using colon and breast cancer atlas datasets to examine their expression patterns. The median expression values of CK2⍺ and CHIP were analyzed in both cancers to determine their roles in tumorigenesis (Fig. 2C). Additionally, expression levels were also examined across breast cancer subtypes, including ER+ (Estrogen Receptor positive), HER2+ (Human Epidermal Growth Factor Receptor 2 positive), and TNBC (Fig. 2D). Furthermore, Broad Institute’s RNA-Seq data of tissue samples were analyzed to compare CK2⍺ and CHIP expressions across tissues (Fig. 2E). We observed CK2⍺ expression was significantly elevated in COAD and BRCA samples compared to CHIP, confirming their differential expression patterns. Additionally, Hematoxylin and Eosin (H and E) staining and IF analysis of formalin-fixed, paraffin-embedded (FFPE) sections from COAD and BRCA patients (n = 10 each) indicated differential expressions of CK2α and CHIP (Fig. 2F-H). These findings support the negative correlation between CK2⍺ and CHIP in COAD and BRCA.

**Fig. 2.**
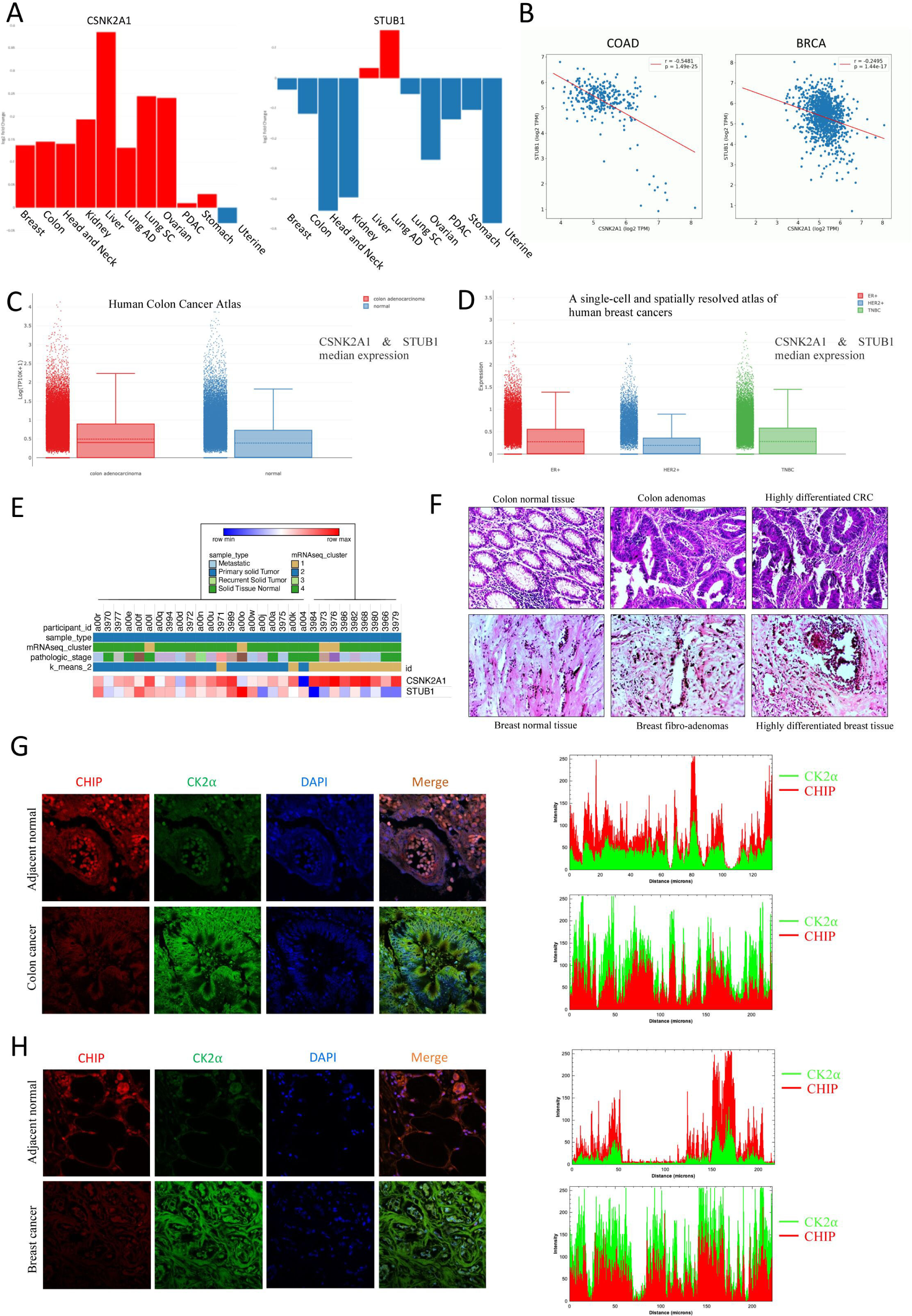
Inverse correlation of CK2α and CHIP expressions in colorectal and breast cancers. **A** CSNK2A1 (CK2α) and STUB1 (CHIP) expression across cancer datasets from cProSite. **B** OncoDB and UALCAN analyses showing significant negative correlation between CK2α and CHIP protein levels in colon adenocarcinoma (COAD, r = –0.54) and breast cancer (BRCA, r = –0.24). **C** Single-cell transcriptomic analysis of colon and breast cancer atlases showing distinct expression patterns of CK2α and CHIP. **D** Expression analysis across breast cancer subtypes (ER+, HER2+, TNBC). **E** RNA-seq tissue data highlighting elevated CK2α expression relative to CHIP in COAD and BRCA. **F-H** H&E staining and immunofluorescence of FFPE sections from COAD and BRCA patient samples (n = 10 each).

### CK2α regulates cellular threshold, localization and tumor suppressive potential of CHIP

The ATP/GTP-binding domain of CK2α enables substrate’s phosphorylation, while the activation segment and P+1 loop regulate its enzymatic activity. The active site is critical for kinase function, making it a target for inhibitors such as TBCA, highlighting CK2α’s importance in cancer cell survival (Fig. 3A, Supplementary Fig. 3A, B) [23–27]. To elucidate CK2α’s role in maintaining CHIP and tumorigenic potential of cancer cells, we successfully inhibited CK2α by either TBCA or, DMAT in HEK293 and cancer cell lines HCT116, and MDAMB-468, followed by IB analysis. The results demonstrated a dose-dependent increase in CHIP protein levels with increasing TBCA and DMAT concentrations (10-40 µM), suggesting that CK2α inhibition directly affects CHIP’s stability (Fig. 3B, Supplementary Fig. 3C). CHIP expression was further examined in a panel of cancer cell lines *viz.*, HCT116, HT29, COS7, 4T1, MDAMB-468, MDAMB-231, and HEK293 after treatment with 20 µM TBCA, and differential expression of CHIP was observed (Fig. 3C, Supplementary Fig. 3D). Next, the intracellular distribution of CHIP and CK2α following TBCA treatment was monitored using IF microscopy (Fig. 3D). Comparative imaging of HEK293, HCT116, MDAMB-468 and MDAMB-231 cell lines showed that TBCA treatment (20 µM for 4 h) altered their cellular levels and localization pattern. TBCA increased CHIP’s nuclear localization. CK2α-mediated compartmentalization of CHIP is critical in cancer cell signalling and tumorigenicity. IB analysis of HCT116 and MDAMB-231 cells treated with DMSO, TBCA, and sorbitol revealed changes in cellular level of CHIP due to altered CK2α activity (Fig. 3E). Next, the dynamic regulation of CHIP following CK2α overexpression was assessed through a 12 h time-course experiment in HEK293 and MDAMB-468 cells (Fig. 3F). IB analysis showed fluctuations in CHIP levels, suggesting CK2α’s role in PTM of CHIP and its stability. Next, to assess the effects of CK2α inhibition on tumor cells, 3D spheroid culture assay was conducted using 4T1 (mouse breast cancer) and HCT116 cells. The spheroids were treated with TBCA for 48 h, followed by a CFDA-SE live-dead assay was performed (Fig. 3G-H, Supplementary Fig. 3E, F). The results showed nice colocalization of CHIP and CK2α in spheroids, changes in the size and integrity of spheroids as well as ratio of live-dead cells, indicating involvement of CK2α in CHIP protein regulation for enhanced tumorigenic potential and growth. Actin was kept as loading control in all the IB analysis.

**Fig. 3.**
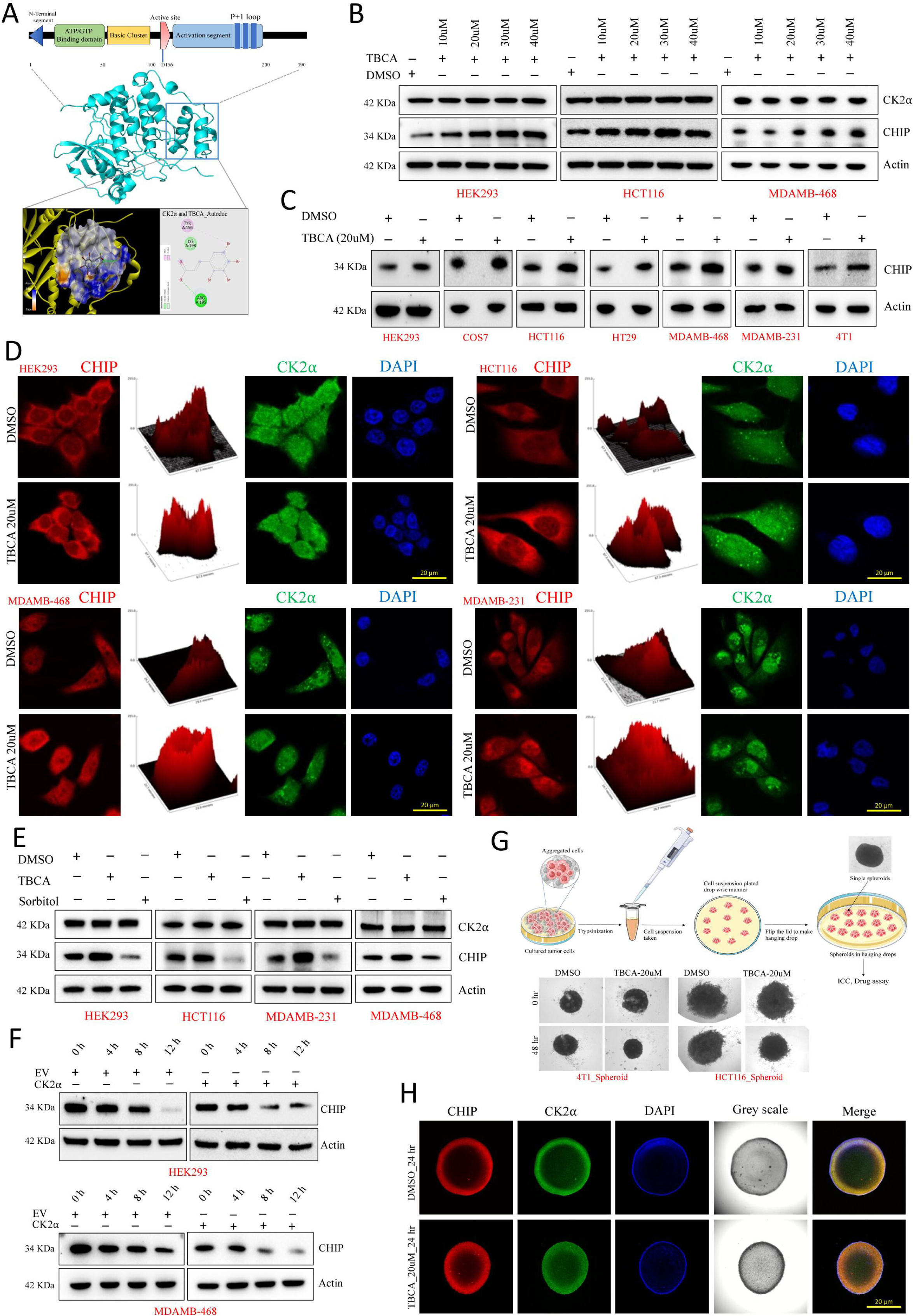
Pharmacological inhibition of CK2α modulates expression, localization, and tumorigenic potential of CHIP. **A** Schematic diagram showing CK2α’s structural domains and -binding sites of its inhibitor. **B** IB analysis of HEK293, HCT116, and MDA-MB-468 cells treated with TBCA (10-40 µM) demonstrating dose-dependent stabilization of CHIP. **C** Differential CHIP expression in multiple cancer cell lines after TBCA treatment (20 µM). **D** IF analysis showing altered localization of CHIP and CK2α upon TBCA treatment in HCT116, HEK293, MDA-MB-468, and MDA-MB-231 cells. **E** IB of HCT116 and MDA-MB-231 cells treated with DMSO, TBCA, or sorbitol showing altered CHIP stability and CK2α activity. **F** Time-course analysis of CK2α overexpression in HEK293 and MDA-MB-468 cells showing dynamic regulation of CHIP protein levels. **G-H** 3D spheroid assays with 4T1 and HCT116 cells treated with TBCA showing reduced spheroid integrity and altered cell viability.

### CK2α regulates CHIP’s stability

To assess the reciprocal regulation of CK2α and CHIP, HEK293 cells were transfected with expression plasmid of either CHIP (0.5, 1.0, and 2.0 µg) or CK2α (0.5, 1.0, and 2.0 µg). CHIP overexpression did not affect CK2α levels, whereas CK2α overexpression induced a dose-dependent reduction in CHIP levels, indicating a negative regulation of CHIP’s stability by CK2α (Fig. 4A, Supplementary Fig. 4A). To assess CK2α’s role in CHIP stability *via* PTM, a cycloheximide (CHX) chase assay was conducted in control and CK2α knocked down (using CK2α siRNA) HEK293 cells. Cells were treated with CHX to inhibit protein synthesis, and CHIP degradation was monitored periodically (0, 4, 8, and 12 h). IB showed less CHIP degradation in CK2α-depleted cells, underscoring CK2α’s role in promoting CHIP degradation (Fig. 4B). Similarly, we examined the effect of overexpression and knockdown CK2α in HCT116, MDA-MB-231, MDA-MB-468, and HEK293 cells. We observed a significant reduction and accumulation of CHIP, respectively (Fig. 4C-D, Supplementary Fig. 4B). To determine the subcellular localization of CHIP upon CK2α overexpression, IF imaging was conducted in MDA-MB-468, HCT116, and 4T1 cancer cells. IF showed a decrease in fluorescence intensity of CHIP upon CK2α overexpression, while CK2α depletion increased CHIP levels (Fig. 4E-F). Pearson correlation analysis (R > 0.7) further confirmed a robust inverse correlation between CK2α and CHIP. To examine CHIP regulation by CK2α, HEK293, MDA-MB-231, and MDA-MB-468 cells were transfected with an empty vector (EV), GFP-tagged wild-type CK2α (CK2α-WT), or kinase-inactive mutant (CK2α-K68M), followed by IB analysis. The results showed that CK2α overexpression decreased CHIP protein levels, while CK2α-K68M failed to induce this effect, indicating that CK2α kinase activity is required for CHIP downregulation (Fig. 4G). To further validate the role of CK2α in CHIP regulation, a rescue experiment was designed and performed in HEK293 and MDA-MB-468 cells, wherein cells were treated with TBCA following CK2α overexpression. IB and IF analyses revealed that TBCA treatment on the CK2α overexpressing condition led to a significant increase in CHIP level, indicating that CK2α kinase activity is essential for promoting CHIP degradation most likely through phosphorylation-dependent mechanism (Fig. 4H, Supplementary Fig. 4C). Actin was kept as loading control in all the IB analysis. All these findings support mechanistic insights into the post-translational regulation of CHIP by CK2α in cancer.

**Fig. 4.**
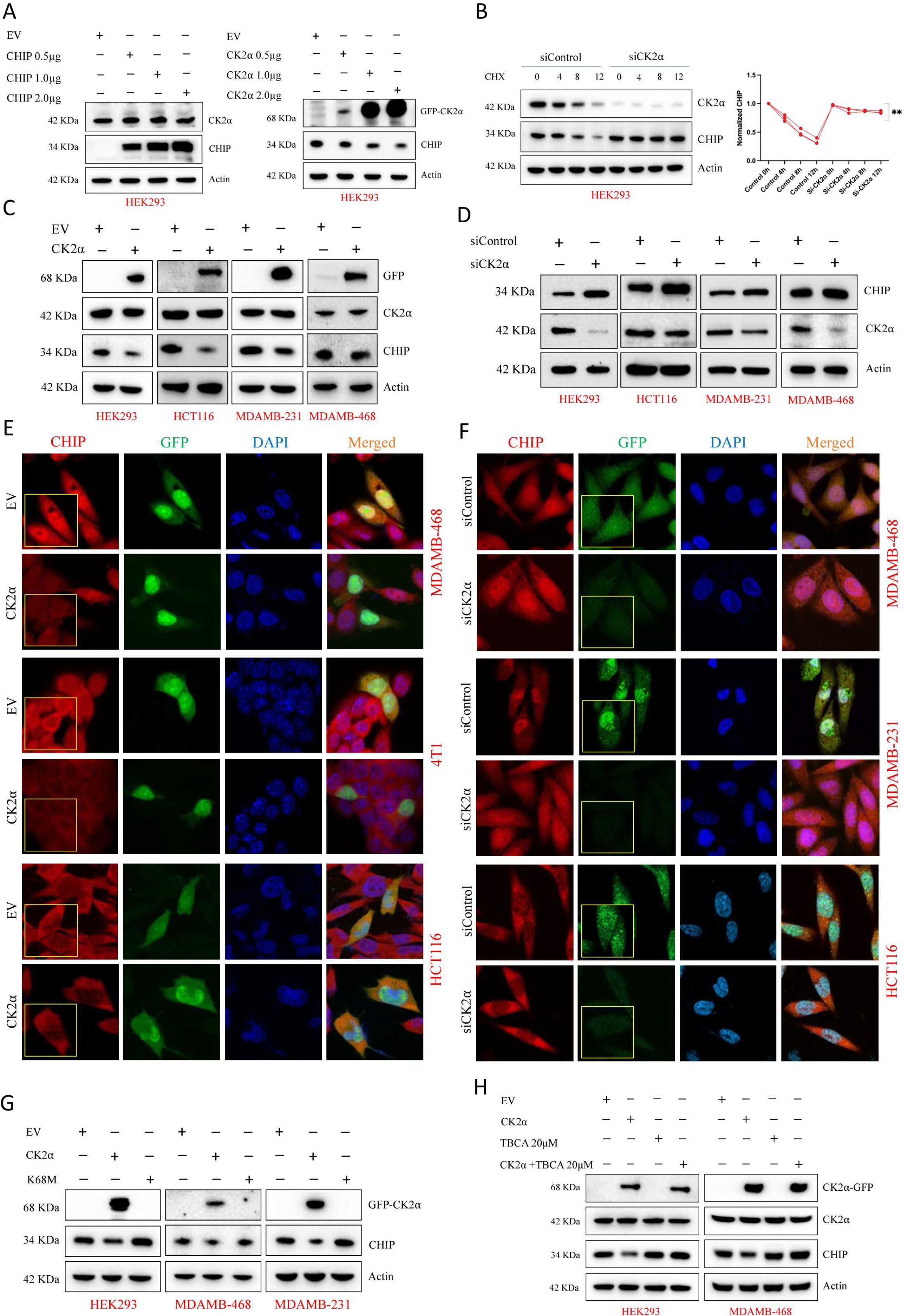
CK2α negatively regulates CHIP’s stability through phosphorylation-dependent degradation. **A** Overexpression of CHIP or CK2α in HEK293 cells showing that CK2α, induces dose-dependent reduction of CHIP protein level. **B** Cycloheximide (CHX) chase assay in HEK293 cells with CK2α knockdown showing reduced CHIP degradation. **C-D** IB analysis in HEK293, HCT116, MDA-MB-231, and MDA-MB-468 cells showing downregulation of CHIP upon CK2α overexpression and accumulation. **E-F** IF analysis in MDA-MB-468, HCT116, 4T1, and MDA-MB-231 cells showing reduced fluorescence of CHIP with CK2α overexpression and increased when depleted; Cells were stained for CHIP (red) and CK2α (green), and nuclei were counterstained with DAPI (blue). Pearson’s correlation confirmed inverse association. **G** IB of cells transfected with CK2α-WT or kinase-deficient mutant CK2α-K68M showing that kinase activity is required for CHIP downregulation. **H** Rescue experiment in HEK293 and MDA-MB-468 cells: TBCA treatment restored CHIP levels despite CK2α overexpression. The graphs were represented as mean ± S.D., n = 3, Significance levels *p≤ 0.05, **p≤ 0.01 and ***p≤ 0.001.

### CK2α promotes ubiquitination and degradation of CHIP

To delineate the molecular mechanism by which CK2α regulates CHIP’s stability, we investigated whether phosphorylation by CK2α influences CHIP’s ubiquitination and subsequent proteasomal degradation. A schematic diagram illustrating the constructs with various domain [28] mutations representing CHIP’s domain-specific function (Fig. 5A). To determine CK2α’s role in CHIP stability, HEK293 cells were co-transfected with CHIP-WT or CHIP mutants *viz.*, K30A (TPR mutant), H260Q (E3 ligase-deficient mutant), ΔUbox (U-box deletion mutant), and TPR along with GFP-tagged CK2α. 48 h post-transfection, IB analysis was performed, results showed decreased CHIP-WT level, indicating that CK2α negatively regulates CHIP stability. The K30A mutant, which disrupts CHIP’s interaction with Hsp70/Hsp90, shows partial resistance to CK2α-induced degradation. The H260Q and ΔUbox mutants impair CHIP ubiquitin ligase activity, displayed increased stability compared to CHIP-WT, indicating that CHIP’s auto-ubiquitination is necessary for CK2α-directed degradation (Fig. 5B-C). The graphical illustration showing CK2α-mediated phosphorylation on CHIP leading to proteasome-mediated degradation (Fig. 5D). Next, HCT116 cells overexpressing CK2α were treated with TBCA, followed by IB analysis, results indicating that inhibition of CK2α activity results in reduced ubiquitination and stabilization of CHIP protein. Additionally, IP followed by IB with anti-ubiquitin antibody confirmed that CK2α inhibition suppressed ubiquitination and degradation of CHIP (Fig. 5E, Supplementary Fig. 5A).

**Fig. 5.**
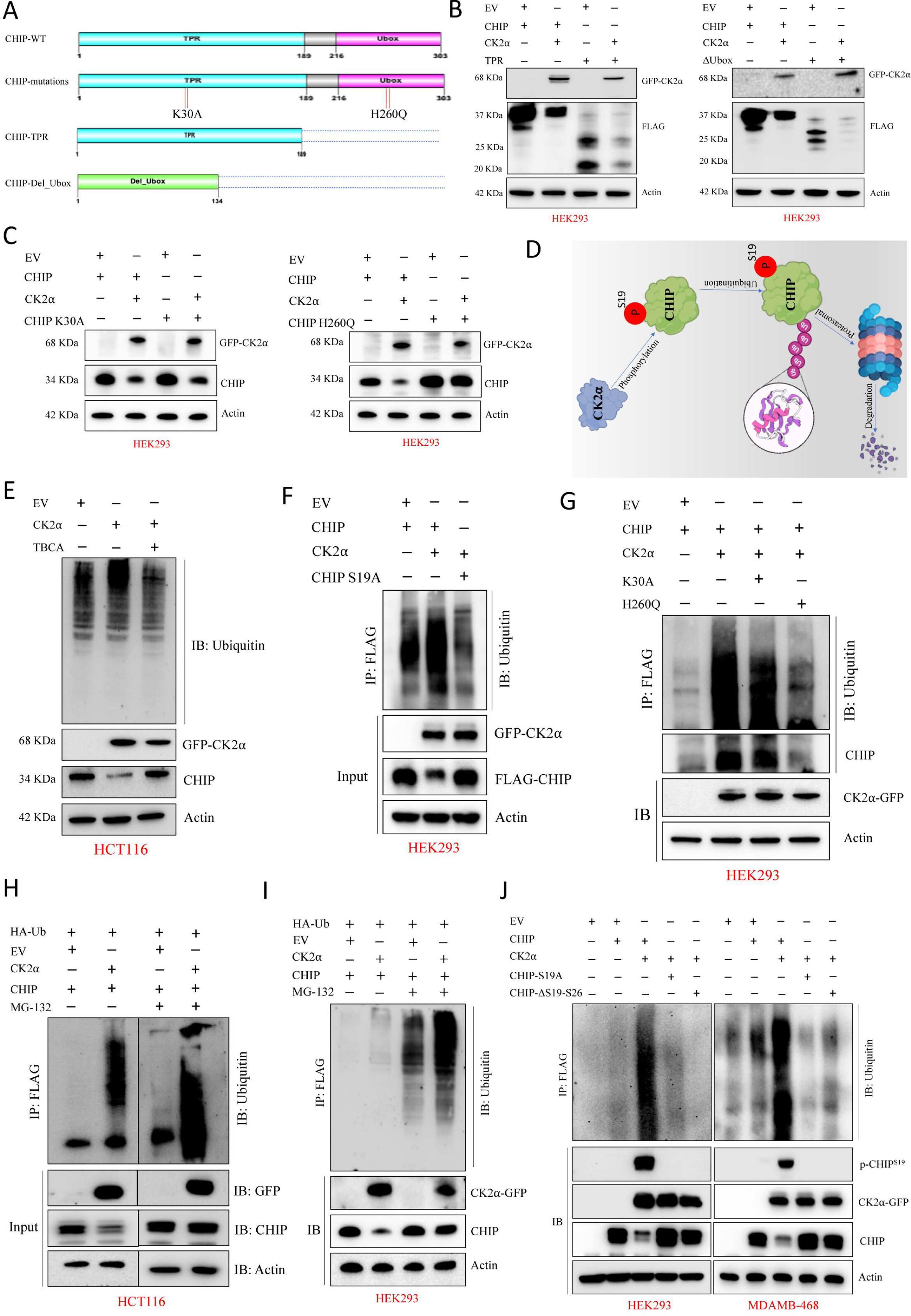
CK2α-mediated phosphorylation of CHIP at S19 promotes its ubiquitination and degradation. **A** Schematic diagram of CHIP constructs with domain-specific mutations. **B-C** IB analysis of HEK293 cells co-transfected with CK2α and CHIP-WT or its mutants (K30A, H260Q, ΔUbox) showing that CK2α reduces CHIP-WT levels, while mutants impairing chaperone binding or E3 ligase activity exhibit increased stability. **D** Graphical representation of CK2α-mediated phosphorylation leading to proteasome-dependent CHIP degradation. **E** IB and IP analysis in HCT116 cells showing that CK2α inhibition by TBCA stabilizes CHIP due to reduced ubiquitination. **F** IP and IB analysis of CHIP-WT and phosphorylation-resistant mutant CHIP-S19A co-transfected with ubiquitin, showing reduced ubiquitination in CHIP-S19A. **G** IP-IB analysis confirming the role of the TPR domain (K30A mutant) in regulating CHIP ubiquitination. **H-I** Co-transfection of CK2α, CHIP, and ubiquitin in HCT116 cells showing that CK2α enhances CHIP ubiquitination, which is reduced upon inhibition. **J** IP followed by IB in HEK293 and MDA-MB-468 cells demonstrating that CK2α overexpression promotes CHIP ubiquitination, whereas CHIP-S19A mutant shows impaired ubiquitination.

Next, to determine whether CK2α-dependent phosphorylation is necessary for CHIP ubiquitination, we generated and used a phosphorylation-resistant CHIP mutant (CHIP-S19A). HEK293 cells were co-transfected with either FLAG-tagged CHIP-WT or CHIP-S19A along with HA-tagged ubiquitin. 48 h post-transfection, CHIP was immunoprecipitated using an anti-FLAG antibody, and ubiquitination level was assessed by IB with an anti-HA antibody (Fig. 5F). IB showed reduced ubiquitination in the CHIP-S19A mutant compared to CHIP-WT, indicating the importance of S19 phosphorylation for CHIP ubiquitination. The decreased ubiquitin conjugation in the CHIP-S19A demonstrates the role of CK2α-mediated phosphorylation in CHIP turnover. This indicates that CK2α-dependent S19 phosphorylation is necessary for ubiquitination and proteasomal degradation of CHIP. Further, HEK293 cells were co-transfected with CK2α and CHIP-WT or K30A/H260Q mutant, followed by IP and IB. Our results confirmed that K30A domain of CHIP plays major role in its ubiquitination (Fig. 5G).

Additionally, HCT116 cells were co-transfected with FLAG-tagged CHIP, HA-tagged ubiquitin, and GFP-tagged CK2α. 42h post-transfection, cells were treated with MG132, followed by IP using anti-FLAG antibody and IB with anti-ubiquitin antibody showed increased ubiquitin conjugation to CHIP, indicating that CK2α enhances CHIP ubiquitination. With CK2α overexpression, ubiquitination of CHIP is increased, showing that CK2α activity affects CHIP’s post-translational fate. These results suggest that CK2α-dependent phosphorylation of CHIP may prime its ubiquitination, affecting protein stability. Similar effect was monitored in HEK293 cells (Fig. 5H-I, Supplementary Fig. 5B). IP assays in MDA-MB-468 and HEK293 cells confirmed CK2α’s role in promoting CHIP ubiquitination at enhanced level. To examine the requirement for serine 19 phosphorylation, cells were transfected with CK2α or the CHIP-S19A mutant, followed by IP and IB of CHIP. The results showed that CK2α overexpression increased CHIP ubiquitination, while the CHIP-S19A mutant showed reduced ubiquitination, highlighting the importance of phosphorylation at this residue (Fig. 5J). These findings strongly suggest that CK2α-mediated phosphorylation of CHIP at S19 serves as a critical regulatory mechanism that facilitates its ubiquitination, thereby accelerating its subsequent recognition and degradation *via* the proteasomal pathway. Actin was kept as loading control in all the IB analysis.

### CK2α restores AKT to promote oncogenesis by phosphorylation dependent degradation of CHIP

Our schematic model illustrates that CHIP, an E3 ubiquitin ligase, targets AKT for ubiquitination and proteasomal degradation, thereby reducing its stability and active phosphorylation level (Fig. 6A). Analysis of TNBC patient-derived xenograft (PDX) proteomes showed an inverse correlation between CHIP and AKT, indicating a potential regulatory relationship between these two proteins (Fig. 6B). Compared to CK2α overexpression in MDAMB-468, CK2α and CHIP in HCT116 and HEK293 shows increase in phospho-AKT (Ser473 and Ser129) (Supplementary Fig. 6A, B). IB analysis was performed in HEK293 cells after overexpression of CHIP which revealed a marked decrease in total AKT and phospho-AKT (Ser473 and Ser129), suggesting that CHIP negatively regulates AKT and results in substantial reduction of its activation (Fig. 6C). Alternatively, knockdown of CHIP using siRNA (siCHIP) in HEK293 cells resulted in elevated levels of AKT and pAKT (Ser473 and Ser129), supporting negative regulatory function of CHIP on AKT stability and activation (Fig. 6D). Additionally, co-transfection of CK2α with CHIP rescued AKT from CHIP-mediated degradation and restored its active phosphorylations (Fig. 6E). Moreover, IF imaging showed that CK2α overexpression led to increased AKT phosphorylations at both Ser129 and Ser473 in HCT116 and HEK293 cells (Fig. 6F, Supplementary Fig. 6C, D).

**Fig. 6.**
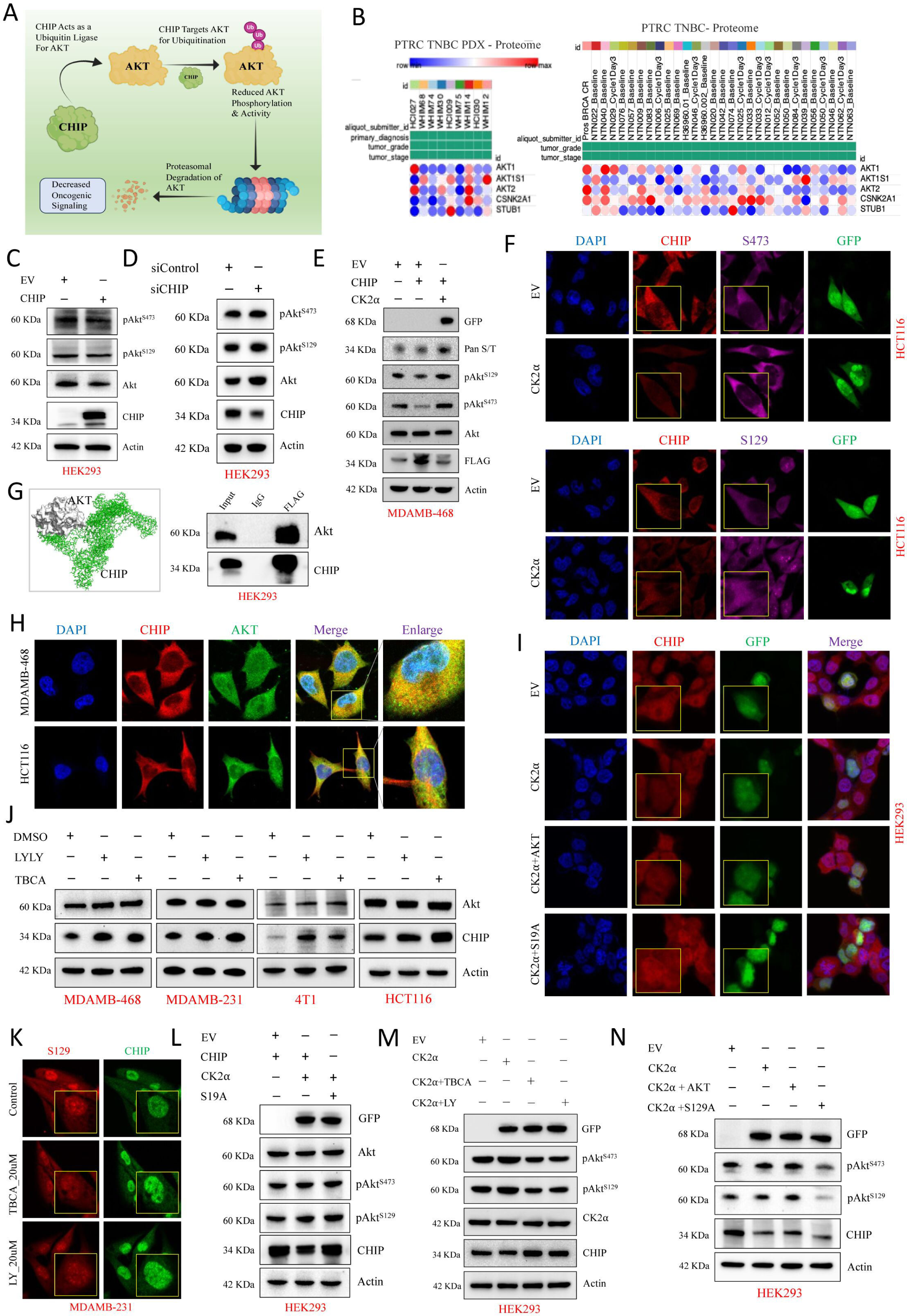
CK2α-CHIP-AKT axis regulates AKT stability and oncogenic signalling. **A** Schematic model showing CHIP-mediated ubiquitination and degradation of AKT. **B** IB of TNBC PDX proteomes showing inverse correlation between CHIP and AKT expressions. **C-D** Overexpression of CHIP reduces, whereas CHIP knockdown increases, AKT and pAKT (Ser473, Ser129) levels in HEK293, HCT116, and MDAMB-231 cells. **E** Co-expression studies demonstrate CK2α rescues AKT from CHIP-mediated degradation and enhances pAKT. **F** IF imaging confirms increased pAKT (Ser129, Ser473) upon CK2α overexpression. **G-H** Docking, co-IP, and colocalization assays reveal direct CHIP-AKT interaction. **I-K** CHIP-S19A mutant resists CK2α- and AKT-mediated downregulation; pharmacological inhibition of CK2α or AKT restores CHIP levels across TNBC cell lines. **L-M** CK2α promotes CHIP degradation, whereas its inhibition rescues CHIP and reduces pAKT. **N** Co-expression with AKT-S129A indicates partial involvement of AKT in CK2α-mediated CHIP degradation.

Next, we performed molecular docking of AKT (PDB: 10CL) with CHIP (PDB: 2C2L) using ClusPro server to assess their interactions and IP assay to further confirm their physical interaction, which supports direct regulatory control of CHIP on AKT (Fig. 6G, Supplementary Fig. 6E). We also observed their colocalization in MDAMB-468 and HCT116 cells by IF study (Fig. 6H). Additionally, overexpression of CHIP-S19A rescued the CHIP level from either CK2α or a combination of CK2α and AKT-mediated CHIP downregulation, indicating that S19 is the major site for phosphorylation (Fig. 6I). To further strengthen our data, we have shown restoration of CHIP levels upon inactivation of either CK2α or AKT by using their respective inhibitors TBCA and LY-294002, in MDAMB-231 and MDAMB-468 (TNBC) as well as 4T1 and HCT116 cell lines, indicating a conserved mechanism across different cancer cell lines by WB and IF analyses (Fig. 6J-K, Supplementary Fig. 6F).

Next, HEK293 cells were co-transfected with either CHIP-WT or CHIP-S19A and CK2α, where EV-transfected cells was kept as control. WB analysis revealed that CK2α led to a decrease in CHIP levels, while the CHIP-S19A attenuated this effect, indicating that CK2α promotes CHIP degradation, through its S19 phosphorylation (Fig. 6L). Additionally, pharmacological inhibition of either CK2α or AKT restored CHIP levels in CK2α-overexpressing HEK293 cells (Fig. 6M). Here, we observed that the reduced phosphorylation of AKT at both S129 and S473 is due to the inhibition of either CK2α or AKT, suggesting that degradation of CHIP occurs by direct involvement of CK2α and additionally through “CK2α-AKT axis”. To further dissect the mechanistic link [29], cells were co-transfected with CK2α and either AKT-WT or AKT-S129A in HEK293 cells. Our IB results indicate that AKT is partially involved in CK2α mediated degradation of CHIP (Fig. 6N). Actin was kept as loading control in all the IB analysis. Altogether, CHIP promotes AKT degradation and suppresses oncogenic signalling, whereas CK2α-mediated phosphorylation of CHIP accelerates its degradation and stabilizes AKT. This CK2α-CHIP-AKT axis diminishes CHIP’s regulatory effect on AKT, facilitating its oncogenic potential.

### Functional validations of CK2α-CHIP-AKT axis *in vitro*

To assess CK2α-mediated phosphorylation of CHIP, migration assays were performed in HEK293 and HCT116 cells, after overexpression of CK2α, CHIP or CHIP-S19A either alone or in combination as depicted. Here, CHIP reduced cell migration, a tumor suppressive action, while CK2α co-expression reversed this effect. The CHIP-S19A mutant have reduce the effect despite CK2α co-expression, confirming CHIP-S19 phosphorylation was necessary to restores migration. These results showed CHIP acts as a cell migration regulator, modulated by CK2α through S19 phosphorylation, linking proteostasis and cancer cell motility (Fig. 7A). Additionally, we investigated the effect of CK2α-mediated CHIP phosphorylation in tumor cell growth using clonogenic assays in MDA-MB-468 and MDA-MB-231 TNBC cell lines. TBCA treatment or CHIP overexpression reduced the colony formation, showing both CK2α inhibition and elevated CHIP levels suppressed survival of cancer cells. Co-expression of CK2α along with CHIP restored colony formation, demonstrating CK2α counteracts CHIP’s tumor-suppressive function. CHIP-S19A decreased the colony formation ability, confirming phosphorylation at serine 19 was essential for CK2α-mediated reversal (Fig. 7B). These results clearly show that CK2α mediated phosphorylation of CHIP modulates its anti-tumorigenic activity, revealing a signalling axis that balancing proteostasis and cell survival in cancer. To assess phosphorylation of CHIP in cell motility, we conducted wound healing assay in MDA-MB-468 cells with overexpression of CHIP-WT or CHIP-S19A mutant and CK2α, alone and in combination. Cells co-expressing CHIP with CK2α showed increased wound closure over 48 h, while CHIP-S19A showed delayed motility, indicating CHIP’s anti-migratory effect depends on CHIP’s E3 ligase activity, but its S19 phosphorylation by CK2α inhibiting this function (Fig. 7C). We also examined that overexpression of CHIP-S19A rescued eliminated CHIP’s apoptotic function by CK2α, showing S19 phosphorylation is crucial to regulate CHIP’s activity. Moreover, in HEK293 cells, overexpression of CHIP increased the levels of pro-apoptotic proteins *viz*., Bax, Bim, and caspase-3, while CK2α co-expression suppressed these apoptotic markers. Deletion of CHIP region S19-S26 abolished the effect of CHIP, confirming N-terminal part of CHIP is important for phosphorylation driven role. Additionally, kinase-dead CK2α mutant failed to inhibit CHIP-mediated apoptosis, showing CK2α’s kinase activity is important for oncogenesis through modulation of CHIP (Fig. 7D).

**Fig. 7.**
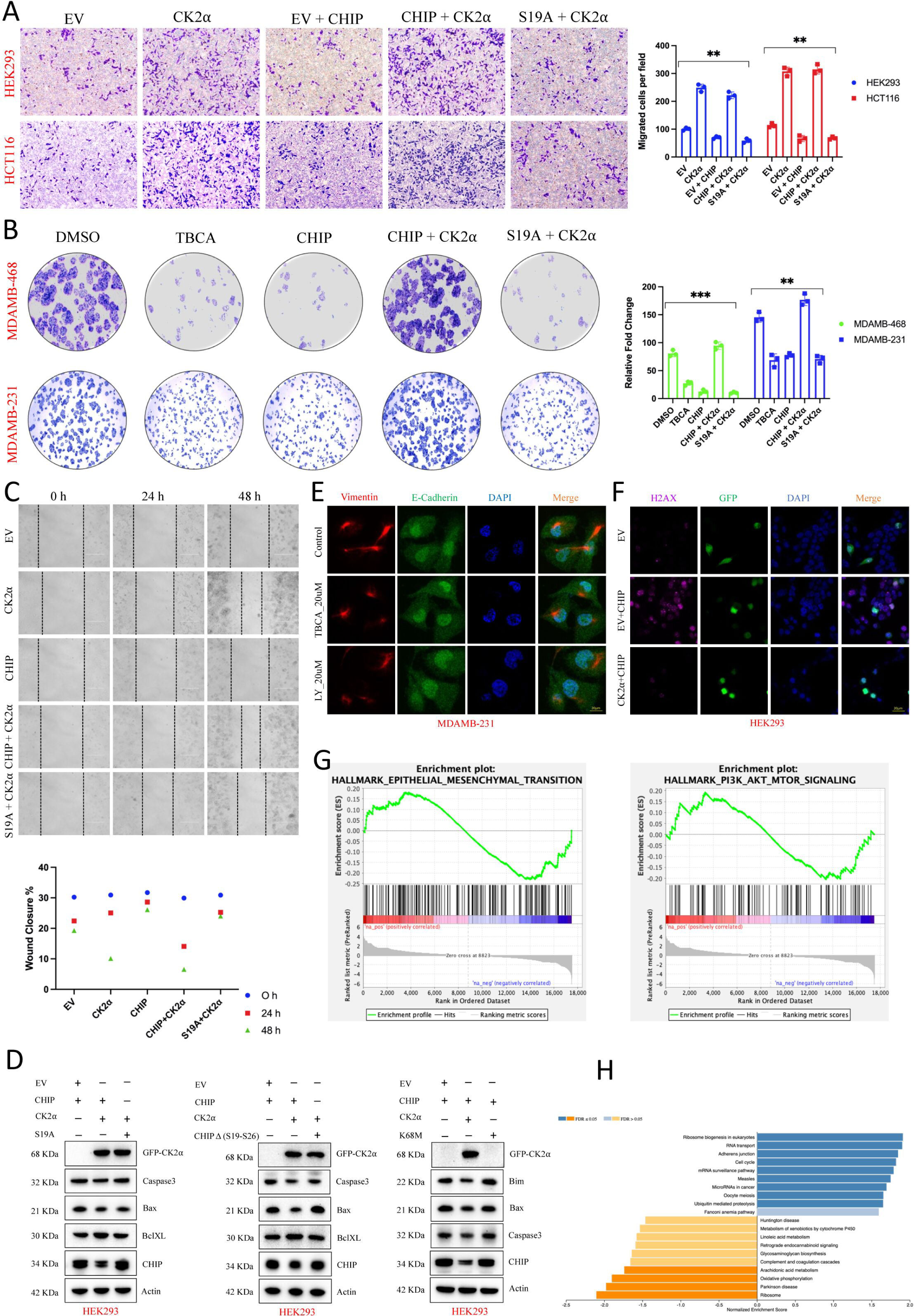
Phosphorylation of CHIP by CK2α regulates apoptosis, migration, and oncogenic potentials. **A** Wound-healing and migration assays in HEK293 and HCT116 cells show CHIP suppresses cell migration, while CK2α co-expression reverses this effect; CHIP-S19A mutant retains anti-migratory activity. **B** Clonogenic assays in TNBC cells reveal that CHIP or CK2α inhibition reduces colony formation, whereas CK2α co-expression rescues this effect; CHIP-S19A fails to restore growth. **C** Wound healing assays confirm CHIP suppresses motility in MDA-MB-468 cells, which is negated by CK2α *via* its S19 phosphorylation. **D** CHIP enhances pro-apoptotic markers (viz., Bax, Bim, caspase-3), which are suppressed by CK2α; CHIP-S19A or CK2α-K68M mutant confirm phosphorylation-dependent regulation. **E** IF analysis of MDA-MB-231 cells treated with LY294002 or TBCA shows EMT marker redistribution (viz., E-cadherin, vimentin). **F** γH2AX staining demonstrates co-expression of CHIP and CK2α modulates DNA damage signaling in HEK293 cells. **G** RNA-seq and GSEA analyses identify EMT, PI3K/AKT signaling, and ubiquitin-mediated proteolysis as major pathways altered by CK2α inhibition. **H** TCGA-BRCA transcriptomic analysis links CSNK2A1(CK2α) expression to phosphorylation and ubiquitin-mediated proteolysis. The graphs were represented as mean ± S.D., n = 3, Significance levels *p≤ 0.05, **p≤ 0.01 and ***p≤ 0.001.

To further examine the effects of PI3K-AKT signaling and CK2α inhibition on epithelial-mesenchymal transition (EMT), IF analysis was conducted on MDA-MB-231 cells treated with either LY294002 (20 µM) or TBCA (20 µM). These treated cells were then immunostained for vimentin (mesenchymal marker protein, red) and E-cadherin (epithelial marker protein, green), as well as DAPI (blue) for nuclei. Merged IF images showed changes in distribution and expression of EMT markers after inhibition of either PI3K or CK2α (Fig. 7E). Moreover, to examine CHIP and CK2α’s role in DNA damage responses, IF was performed in HEK293 cells transfected with overexpression of CHIP alone or in combination with GFP-CK2α. After treatment, these cells were stained with phosphorylated H2AX (γH2AX, a DNA damage sensor, pink), GFP (green, showing transfection), and DAPI (blue). Merged images showed nuclear γH2AX accumulation and GFP expression, indicating CK2α and CHIP co-expression affects DNA damage signalling (Fig. 7F, Supplementary Fig. 7A). Additionally, RNA sequencing workflow shows TBCA-treated HEK293, HCT116, and HeLa cells underwent RNA extraction followed by Illumina transcriptomic profiling (Supplementary Fig. 7B). Differential gene expression analyses identified transcriptional changes and DEGs. Gene set enrichment analyses (GSEA) revealed major pathways including EMT, PI3/AKT signalling, and ubiquitin-mediated proteolysis (Fig. 7G). An integrative transcriptomic analysis of TCGA-BRCA cohort using Linkedomics database, categorized by pathological stages (Stage I [N=183], Stage II [N=622], Stage III [N=250], and Stage IV [N=20]), was conducted. KEGG pathway enrichment analysis of genes associated with CSNK2A1 (CK2α) identified phosphorylation-related signaling cascades and ubiquitin-mediated proteolysis as the most enriched pathways (Fig. 7H, Supplementary Fig. 7C-F). Actin was kept as loading control in all the IB analysis. Thus, the CK2α-CHIP axis emerges as a pivotal signalling node linking proteostasis, cell survival, and oncogenic progression through coordinated control of EMT, DNA damage responses, and ubiquitin-mediated proteolysis.

### Tumorigenic potential of CK2α through regulation of CHIP is supported by spheroid and animal model

We further investigated CK2α’s role in CHIP stability and tumorigenesis using 3D spheroids and xenograft models. CK2α phosphorylates CHIP at S19, causing CHIP destruction, preventing cell death, and promoting tumor growth through maintenance of activated AKT (pAKT) and PCNA upregulation. CK2α inhibition by TBCA stabilizes CHIP, activates cell death signals, and reduces tumor growth (Fig. 8A). IF imaging showed DMSO-treated control spheroids maintained integrity with defined morphology. In contrast, spheroids exposed to TBCA for 24 and 48 h showed reduced size, irregular borders, and disrupted architecture, indicating decreased tumor-forming capability (Fig. 8B). To evaluate the effect of CK2α inhibition on 3D tumor spheroid viability, cells were stained with CFDA (green, live cells) and PI (red, dead cells). 4T1 breast cancer spheroids treated with DMSO (control) or 20 µM TBCA were imaged at 24 h. Control spheroids showed strong CFDA fluorescence with minimal PI signal, indicating high viability, whereas TBCA-treated spheroids showed reduced CFDA and increased PI staining at 24 to 48 h, demonstrating time-dependent cell death and significant spheroid size reduction (Fig. 8C). The experimental workflow indicates bilateral tumors generated in BALB/c mice through subcutaneous injection of CK2α overexpressing 4T1 breast cancer cells with Matrigel. When tumors reached measurable sizes, animals received therapeutic compound intraperitoneally. Over 24 days, three doses of TBCA (25 mg/kg) were administered on days 16, 20, and 24, followed by sacrifice for assessment (Fig. 8D). Representative images of tumors harvested from control and drug-treated mice on the 28^th^ day after sacrifice and performed BLI imaging. Comparative analysis revealed marked differences in tumor morphology and apparent volume, suggesting a therapeutic response in the TBCA-treated groups (Fig. 8E-F). Control tumors showed a dense and organized structures. In contrast, TBCA-treated tumors had disrupted shape and less cellular density. Control lung tissue sections showed nodule formation due to spreading of cancer cells, while treated lungs maintained normal appearance (Fig. 8G). IF images of primary tumor sections from control and treated groups were analyzed. Sections were stained for CK2α (green) and P-CHIP^S19^ (red) with DAPI (blue) for nuclei. Control tumors showed strong CK2α and low P-CHIP^S19^, while treated tumors displayed reverse patterns, indicating disrupted CK2α-CHIP signalling after TBCA mediated intervention of CK2α (Fig. 8H). Analysis of IF data showed colocalization of CHIP (red) and CK2α (green) within both nucleus and cytoplasm of control tumors. In contrast, tumors treated with TBCA exhibited reduced CK2α intensity and diffuse CHIP distribution, indicating disruption of the CK2α-CHIP axis (Fig. 8I). Co-staining of CHIP (red) and PCNA (green) showed increased PCNA expression is overlapping with CHIP in control samples, indicating active proliferation, whereas PCNA level was decreased in treated tissue sections, reflecting reduced proliferation due to enhanced CHIP stabilization, leading to suppression of tumor growth (Fig. 8J). Furthermore, apoptotic cell death in tumor sections from control and treated groups were assessed by TUNEL staining. We observed the number of TUNEL-positive nuclei is increased in TBCA-treated tissues compared to control group. Thus, increased number of TUNEL-positive cells in treated tumors, supporting the link between CK2α inhibition and CHIP-mediated apoptosis (Fig. 8K). Here, all scale bars are 20 µm (IF) and 100 µm (for H & E).

**Fig. 8.**
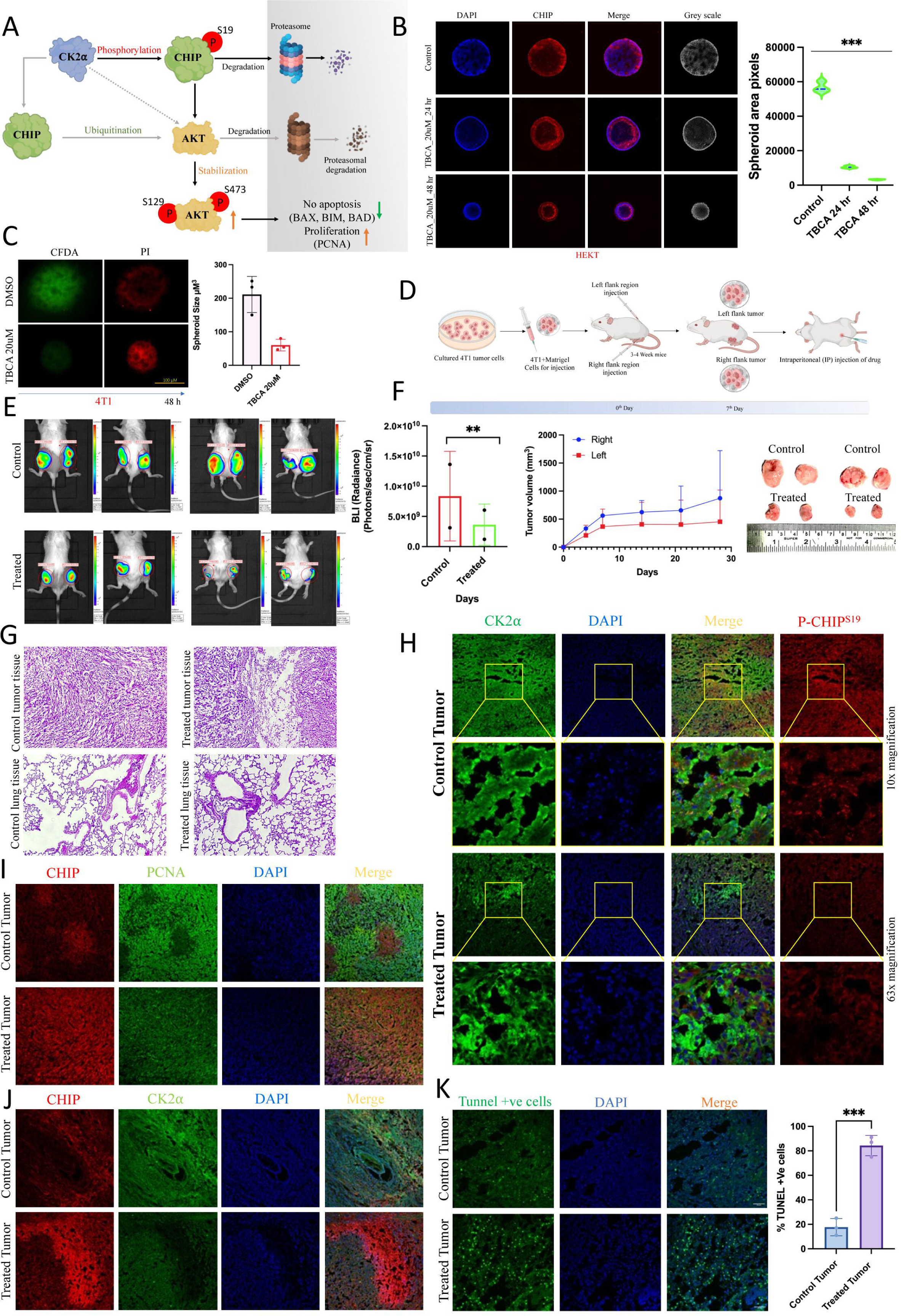
CK2α-mediated CHIP degradation regulates tumor growth and metastasis *in vitro* and *in vivo*. **A** Schematic model illustrating CK2α phosphorylates CHIP at S19, leading to its degradation, pAKT stabilization, and tumor growth, whereas TBCA treatment restores CHIP and promotes apoptosis. **B-C** Confocal and live–dead imaging of 3D spheroids show that TBCA treatment reduces spheroid size, disrupts morphology, increases CHIP expression, and induces apoptosis in a time-dependent manner. **D-F** Experimental design and representative images of bilateral xenograft tumors in BALB/c mice treated with TBCA, showing reduced tumor size and altered morphology by gross observation and BLI imaging. **G** H&E staining of lung sections reveals reduced metastatic spread in TBCA-treated animals. **H-I** IF staining of tumor sections shows altered CK2α and P-CHIP-S19 expression, and reduced colocalization of CHIP and CK2α upon TBCA treatment. **J** CHIP and PCNA co-staining demonstrates decreased proliferation and enhanced CHIP stability in treated tumors. **K** TUNEL assay reveals increased apoptosis in TBCA-treated tumors. The graphs were represented as mean ± S.D., n = 3, Significance levels *p≤ 0.05, **p≤ 0.01 and ***p≤ 0.001.

This study underscores CK2α-CHIP signalling as a mechanistically defined and targetable vulnerability in cancers. Inhibition of CK2α restored CHIP stability and function, inducing apoptosis *in vitro* and tumor regression *in vivo*. These findings highlight the potential of targeting the CK2α-CHIP axis as a therapeutic strategy for solid tumors, represented by our proposed model (Fig. 9).

**Fig. 9.**
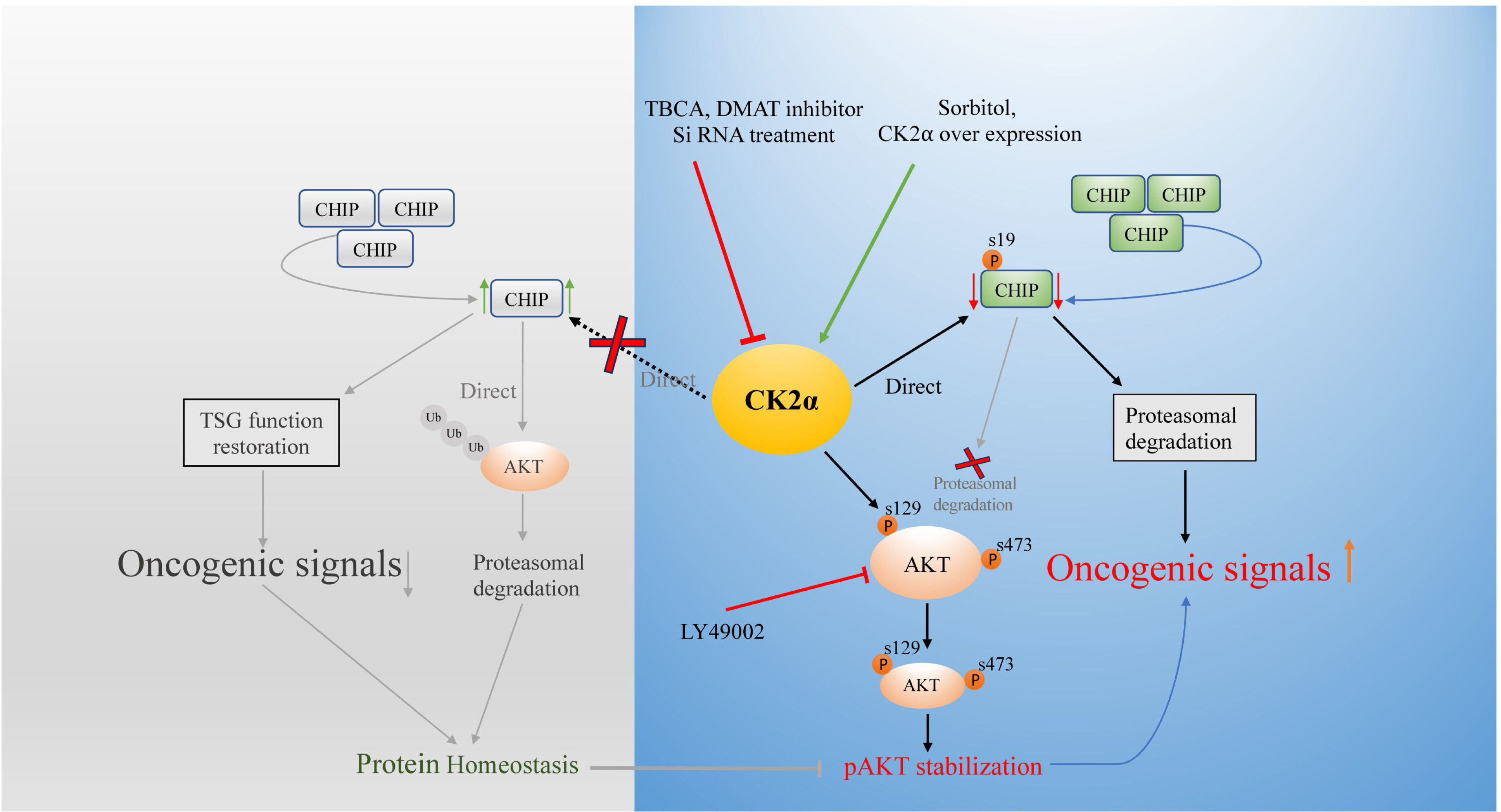
Proposed model illustrating CK2α-mediated regulation of CHIP signalling in oncogenesis. CK2α-driven phosphorylation of CHIP at Ser19 facilitates its proteasomal degradation, thereby attenuating CHIP-mediated ubiquitination of AKT and stabilizing phosphorylated AKT (Ser129/Ser473), culminating in enhanced oncogenic signalling. Conversely, inhibition of CK2α activity (via TBCA, DMAT, or siRNA) restores CHIP integrity, and re-establishing protein homeostasis and tumor-suppressive functions.

## DISCUSSION

CK2α frequently upregulated in many malignancies, regulates E3 ubiquitin ligase CHIP through phosphorylation at serine 19. This kinase affects CHIP’s roles in protein homeostasis, apoptosis, and AKT signalling during tumor progression[1, 30, 31]. We successfully identified CHIP as a CK2α substrate through bioinformatics, mass spectrometry, mutational analyses, and interaction assays. CK2α phosphorylates CHIP’s S19 residue within its N-terminal domain. The interacting interface involved CHIP’s TPR domain, which mediates binding with CK2α. Our data clearly show CK2α phosphorylates CHIP and directs its degradation through ubiquitin tagging, linking kinase signalling to protein homeostasis machinery.

This regulatory mechanism has broad implications in cancer. CK2α-mediated destabilization of CHIP affects protein quality control by reducing ubiquitination of its substrates. As CHIP degrades several oncogenic proteins, its loss under CK2α’s kinase activity may contribute to proteotoxic stress evasion, a cancer hallmark. Inhibition of CK2α using TBCA increased stability of CHIP, indicating kinase activity of CK2α is required for CHIP turnover. Moreover, CHIP-S19A mutation avoids CK2α-mediated phosphorylation dependent degradation, reinforcing this phosphorylation event is important. Additionally, we showed CK2α and CHIP levels has an inverse correlation across transcriptomic and proteomic datasets of colorectal and breast cancer tissues, including bulk and single-cell analyses. IF imaging of tissue sections confirmed their reciprocal expression patterns, particularly in tumor cores CK2α was abundant and CHIP downregulated. Our data suggest CK2α overexpression may serve as a biomarker for loss of CHIP and provide a mechanism for CHIP’s silencing in cancer at post-translational level, independent of transcriptional repression or gene deletion.

To further assess the effect of CHIP depletion in tumor cells, we used two-dimensional and three-dimensional spheroid models. CK2α inhibition restored CHIP and impaired spheroid cohesion and viability, indicating disrupted oncogenic architecture. Also, loss of CHIP is correlated with altered subcellular localization, highlighting its regulation by CK2α activity. Mechanistic analysis revealed CK2α-mediated CHIP degradation affects oncogenic AKT signalling, often dysregulated in cancer. CHIP suppresses oncogenic AKT level and its activity through ubiquitination mediated degradation. Thus, loss of CHIP driven by CK2α results in enhanced AKT phosphorylation at Ser473 and Ser129, amplifying oncogenic signalling. This regulation was validated by using CHIP-S19A and kinase-inactive CK2α-K68M mutants, which restored CHIP stability and reduced AKT level and its activity. Therefore, ’CK2-CHIP-AKT axis’ forms an additional critical node in cell survival, and our findings suggest degradation of CHIP drives CK2α-promoted tumorigenesis.

CHIP suppresses cancer cell migration and clonogenicity. In breast and colon cancer, CHIP’s overexpression inhibited motility and colony formation, this effect is reversed by CK2α co-expression or abolished in presence of phosphorylation-resistant CHIP mutant. These observations showed CK2α acts upstream of CHIP and modulates its anti-migratory and anti-proliferative properties through phosphorylation mediated degradation. CHIP’s E3 ligase activity is required, as loss-of-function due to phosphorylation failed to replicate CHIP(WT)’s tumor-suppressive effects. Additionally, CK2α-mediated CHIP degradation affects stability of its substrates and reprograms cellular behaviour. In mouse xenograft models, CK2α inhibition suppressed tumor growth and metastatic burden due to increased accumulation of CHIP that leads to induced apoptotic signalling. Confocal imaging of treated tumor sections showed reduced colocalization of CK2α and CHIP, diminished expression of proliferation marker (PCNA), and upregulation of apoptotic proteins like Bax, Bim and caspase-3. These results further confirm the CK2α-CHIP relationship is functionally relevant in tumorigenesis, and thus targeting CK2α can restore CHIP activity with therapeutic benefits.

Altogether, our study established a new mechanistic framework linking CK2α activity with stability of tumor suppressor CHIP for downstream oncogenic processes. The identification of a phosphorylation-dependent degradation mechanism of CHIP advances our understanding of how kinases control PTM and the UPS to regulate protein quality control and tumor progression. Moreover, it explains the frequent silencing of CHIP in cancer cells, highlighting its post-translational regulation. From a translational perspective, the reversibility of CHIP degradation through CK2α inhibition offers a novel therapeutic perspective. Here, TBCA, a selective CK2α inhibitor, restores CHIP levels and reactivates its tumor-suppressive functions in multiple cancer models. Thus, CK2α’s druggable nature and specific interaction with CHIP, this axis may be exploited to reinstate proteostasis and restrain hyperactive pathways, such as AKT signalling, in tumors with elevated CK2α expression. Although CK2α activity primarily modulates CHIP stability, other pathways, such as apoptosis and stress-response signalling, may converge on CHIP to fine-tune its function. The interplay between phosphorylation, ubiquitination, and cellular localization determines availability of CHIP within distinct tumor microenvironment (TME). Further investigation of how CK2α cooperates with other kinases or phosphatases to control CHIP turnover could reveal additional regulatory nodes for intervention.

## METHODS

### Inhibitors and knockdown with siRNA

TBCA (CK2α inhibitor, cat.218710) and LY294002 (PI3K/AKT inhibitor, cat.440202) were added at 20 μM for 4 h. To enhance CK2α activity, 0.5 M d-sorbitol was used for 1 h. Cycloheximide (cat.239764; Calbiochem) was administered at 10-25 mM for 4-12 h. MG132 (proteasome inhibitor, cat.474790; Calbiochem) was used at 25 mM for 6 h. Primary antibodies included CK2α, pAKT S129/S473, Caspase-3, Bax, Bim (Cell Signalling), pCHIP S19 (Affinity), CHIP, GFP, AKT-Ub, PCNA (Santa Cruz), CHIP (AbCam), and Actin, GFP, HA-, FLAG-tag (Sigma). HRP-conjugated secondaries for goat, mouse, and rabbit were from Sigma. Scramble siRNA, CK2α (sc-29918), and CHIP siRNA (sc-43555; Santa Cruz Biotechnology) were used at a concentration of 30 nM. A detailed list of antibodies and online tools used in this study were listed in Table S1 and S2.

### Mass spectroscopy

To overexpress CK2α and CHIP, HEK293 cells were transfected with respective plasmids and untransfected cells kept as controls. After 48 h, cell lysates underwent SDS-PAGE, and CHIP-containing gel regions from both conditions (in duplicates) were excised. LC-MS/MS was performed after in-gel trypsin digestion and followed by software based analysis. Peptides (1 μg/injection) were separated using a reverse-phase C18 column on a Thermo Scientific Nano-LC. A gradient of 3-45% Buffer B over 135 min was applied, ramped to 95% at 140 min, held for 10 min, and then re-equilibrated. A 50-minute blank run minimized carryover. Mass spectrometry was conducted using a Q Exactive Orbitrap (Thermo Fisher) as per discussed previously [32]. A detailed list of peptides are incorporated in Table S3.

### Immunoblotting

Cells were lysed in whole-cell lysis buffer as described previously ^23,24^. Forty micrograms of total protein per sample were resolved by SDS-PAGE, transferred to PVDF membranes, and probed with primary antibodies (overnight, 4°C) followed by HRP-conjugated secondary antibodies (Sigma-Aldrich, 2 h, RT). Blots were developed using Classico ECL substrate (Millipore) and visualized with a Bio-Rad ChemiDoc system. Band intensities were quantified using ImageJ, normalized to respective loading controls.

### Site directed mutagenesis

Site-directed mutagenesis (SDM) was performed using PCR as per our previously standardised protocol [33, 34]. Initial PCRs used subcloning primers (with restriction sites) paired with mutagenic primers. Products were gel-extracted, quantified, and mixed (1:1, 100 ng each) for a second PCR using subcloning forward and reverse primers. The final product was purified for ligation and transformation. CHIP mutants (CHIP-S19A and CHIP-Del19-26) were generated from pIRES-CHIP using QuickChange XL SDM Kit (Agilent). Primer sequences are listed in Table S4.

### Ubiquitination assay

Cells were treated with MG-132 for 4-6 h to accumulate polyubiquitinated proteins, then lysed in SDS-lysis buffer after a 10-minute boil. Lysates were diluted 10-fold with IP-Lysis Buffer containing 10 mM NEM and protease inhibitors. Immunoprecipitation was performed with specific antibodies, and HA-tagged proteins were detected using an anti-HA antibody as per our previous protocol [7].

### Animal maintenance and syngeneic flank tumor model

BALB/c mice (3-4 weeks old) were obtained from the IICB animal facility and acclimatized for one week. Early passage 4T1 breast cancer cells stably expressing CK2α (1 × 10^6^) were injected at the flanks using our established protocols. Tumor growth was monitored using Vernier callipers, followed by TBCA treatment (25 mg/kg). Mice were euthanized with urethane, and tumors were collected. Tissues from the control, vehicle, and treated groups (n=3 per group) were fixed in 10% neutral buffered formalin, paraffin-embedded, and sectioned at 5 μm for histology and ICC. H&E staining was performed for histological analysis.

### Immunofluorescence microscopy

Immunofluorescence microscopy of cells was performed as described previously [33, 35, 36]. Primary antibodies for CK2α (CST) and CHIP (Santa Cruz, Abcam) were used with Alexafluor-conjugated secondary antibodies (488, 546, 594, 647). Nuclei were stained with Hoechst 33342 (Life Technologies). Imaging was performed at 63x using a Zeiss LSM 980 confocal microscope with ZENblue3 software; Fiji and ImageJ software was used for analysis.

### Immunohistochemistry (IHC)

Immunohistochemical staining was performed on FFPE sections as previously described [36]. Slides were imaged at 40x magnification using the EVOS XL Cell Imaging System (Life Technologies-Thermo Fisher Scientific). Staining intensity was quantified using the H-score method, calculated by assessing both the percentage of positive cells and staining intensity. Scoring was performed by two independent, blinded observers.

### Wound healing (scratch) and Colony formation assays

The wound healing assay was conducted on HEK293, HCT 116, and other cell lines following the method outlined as described previously [33, 35]. Images were captured 48 h post-transfection. Additionally, colony formation assay was performed on HEK293, HCT 116, MDAMB-231, MDAMB-468, 4T1, and other cells, as previously described [35].

### Spheroid assay

Spheroids were formed with HEK293, HCT116, MDA-MB-468, and 4T1 cell lines using the hanging drop method. Cells (1×10^4^ per 10 µL drop) were placed on the inner side of Petri dish lids, inverted over PBS-filled bases to maintain humidity, and incubated at 37 °C with 5% CO₂ for 72-96 h. The formed spheroids were transferred to low-attachment 96-well U-bottom plates for drug treatment, immunofluorescence, and imaging [37–39].

### Statistical analysis

Statistical significance was determined using the two-sample Student’s t-test, with significance levels indicated as * (P ≤ 0.05), ** (P ≤ 0.01), and *** (P ≤ 0.001). Statistical evaluations were conducted using GraphPad Prism 9 (https://www.graphpad.com/scientific-software/prism).

## DATA AVAILABILITY

Data supporting the findings of this study are available from the corresponding authors upon reasonable request.

## AUTHOR CONTRIBUTIONS

SK & MKG: Conceived the idea, wrote & revised the manuscript. SK: Performed and analyzed data under MKG supervision. MC: helped SK for data generation and manuscript drafting. MB: revised the manuscript and edited accordingly.

## ACKNOWLEDGEMENTS

We would like to acknowledge Dr. Uttara Chatterjee (Park Clinic, Kolkata, India) for providing the patient samples. Additionally, we also acknowledge the Academy of Scientific and Innovative Research, Ghaziabad-201002, India, for the PhD registration of SK and MC.

## ETHICS APPROVAL

All animal procedures were performed under the guidelines of the institutional review board and the ethics committee of CSIR-Indian Institute of Chemical Biology, Kolkata and Park Clinic, Kolkata, India.

## Supplementary Files

### 1. Materials and methods

#### 1.1. Cell culture and transfection

Human colon cancer cell lines HCT116, SW-480, and HT29; breast cancer cell lines MDAMB-231, MDAMB-468, and MCF-7; and human embryonic kidney cell line HEK293 were cultured using a standard protocol (Dulbecco’s Modified Eagle’s Medium with 10% heat-inactivated Fetal Bovine with antibiotics. The mouse cell line 4T1 was cultured per the standard protocol. For transfection, polyethylenimine (PEI, Sigma Aldrich) was used in HEK293 cells, while Lipofectamine 3000 (Invitrogen) was used for other cancer cell lines to achieve 70-85% transfection efficiency. siRNAs were introduced using Lipofectamine RNAiMAX (Invitrogen).

#### 1.2. Plasmid constructs

The plasmids pGZ21dx-GFP-CK2-WT (GFP-CK2), pIRES-CHIP (CHIP-WT), pIRES-CHIP-H260Q, pIRES-CHIP-K30A, and pGZ-AKT1-GFP have been previously documented. PRK5-HA-Ubiquitin-WT was obtained from Addgene.

#### 1.3. Western blot analysis and Immunoprecipitation (IP)

Cells were lysed in Whole-Cell Lysis Buffer, sonicated (30% amplitude, 10s ON/OFF for 30s), and centrifuged at 12,000g for 15 min. Lysates (40–60 µg) were boiled in 2X loading buffer, resolved by SDS-PAGE, and transferred to PVDF membranes. Blots were incubated with primary antibodies overnight at 4 °C, then with secondary antibodies for 2 h at room temperature. Detection was performed using Classico ECL (Millipore) and Bio-Rad Chemi-Unk imaging. For IP, cells were lysed in IP-Lysis buffer on ice (2 h), pre-cleared with Sepharose A/G beads (2 h), and centrifuged. Supernatants were incubated with specific antibodies overnight at 4 °C, followed by bead blocking for 4 h. Beads were washed thrice, boiled in loading buffer, and analyzed by western blotting. VeriBlot HRP reagent (ab131366) minimized IgG background. Inputs (3%) were analyzed separately. Co-IP was conducted in HEK293, HCT116, and MDAMB-468 cells, as described.

**Supplementary Fig. 1.**
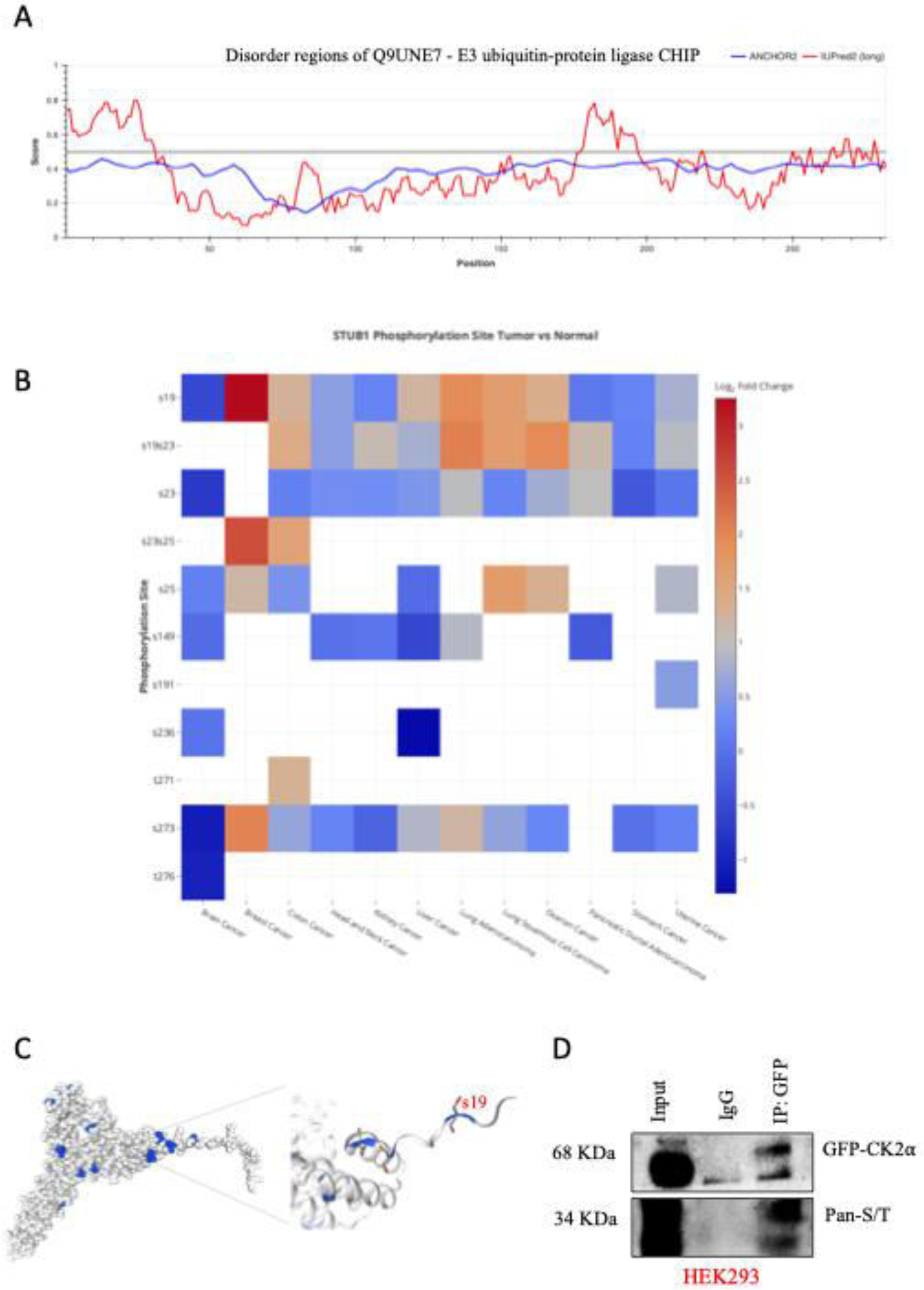
(**A**) Schematic representation of predicted disorder regions within CHIP (UniProt ID: Q9UNE7), highlighting phosphorylation-prone serine/threonine residues. (**B**) *STUB1* predicted phosphorylation residues by cProSite in tumor vs normal. (**C**) Residual visualization of serine 19 (S19) in the N-terminal region of CHIP as a critical CK2α target site. (**D**) Co-immunoprecipitation assay in HEK293 cells expressing CK2α shows that CK2α interacts with CHIP, with input and IgG controls confirming specificity followed by Pan-S/T verification.

**Supplementary Fig. 2.**
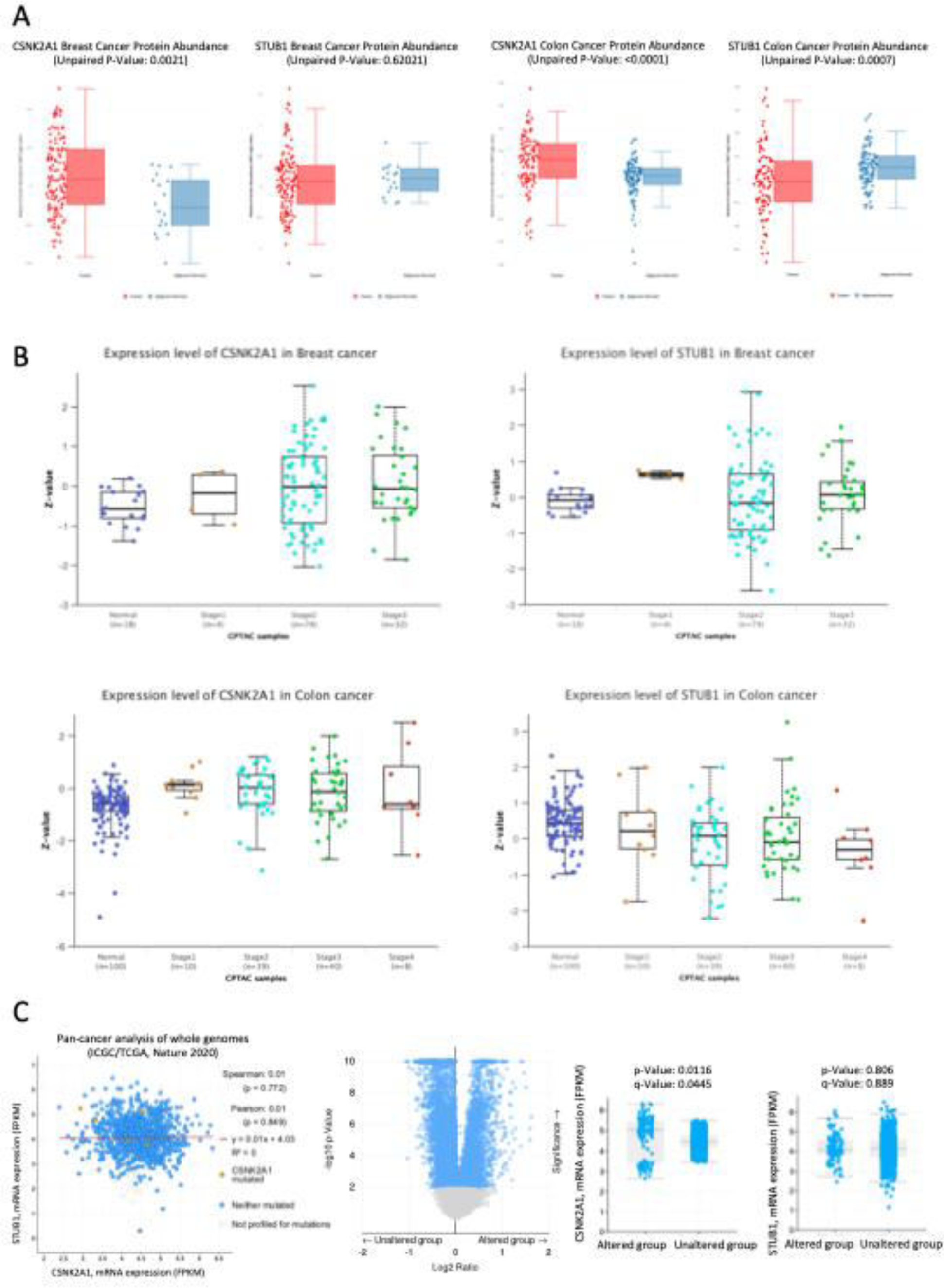
(**A**) Protein abundance analysis in breast and colon cancer tissues shows significantly elevated CSNK2A1 levels (p = 0.0021 in breast; p < 0.0001 in colon), while STUB1 levels remain unchanged in breast cancer (p = 0.62021) but are significantly reduced in colon cancer (p = 0.0007). (**B**) mRNA expression analysis from TCGA indicates higher CSNK2A1 expression in the altered group compared to the unaltered group (p = 0.0116, q = 0.0445), whereas STUB1 mRNA levels show no significant difference between groups (p = 0.806, q = 0.889). (**C**) Pan-cancer analysis of whole-genome data (ICGC/TCGA, Nature 2020) reveals distinct distribution patterns of CSNK2A1 and STUB1 expression across altered and unaltered tumor groups, further supporting a divergence in their roles during tumorigenesis.

**Supplementary Fig. 3.**
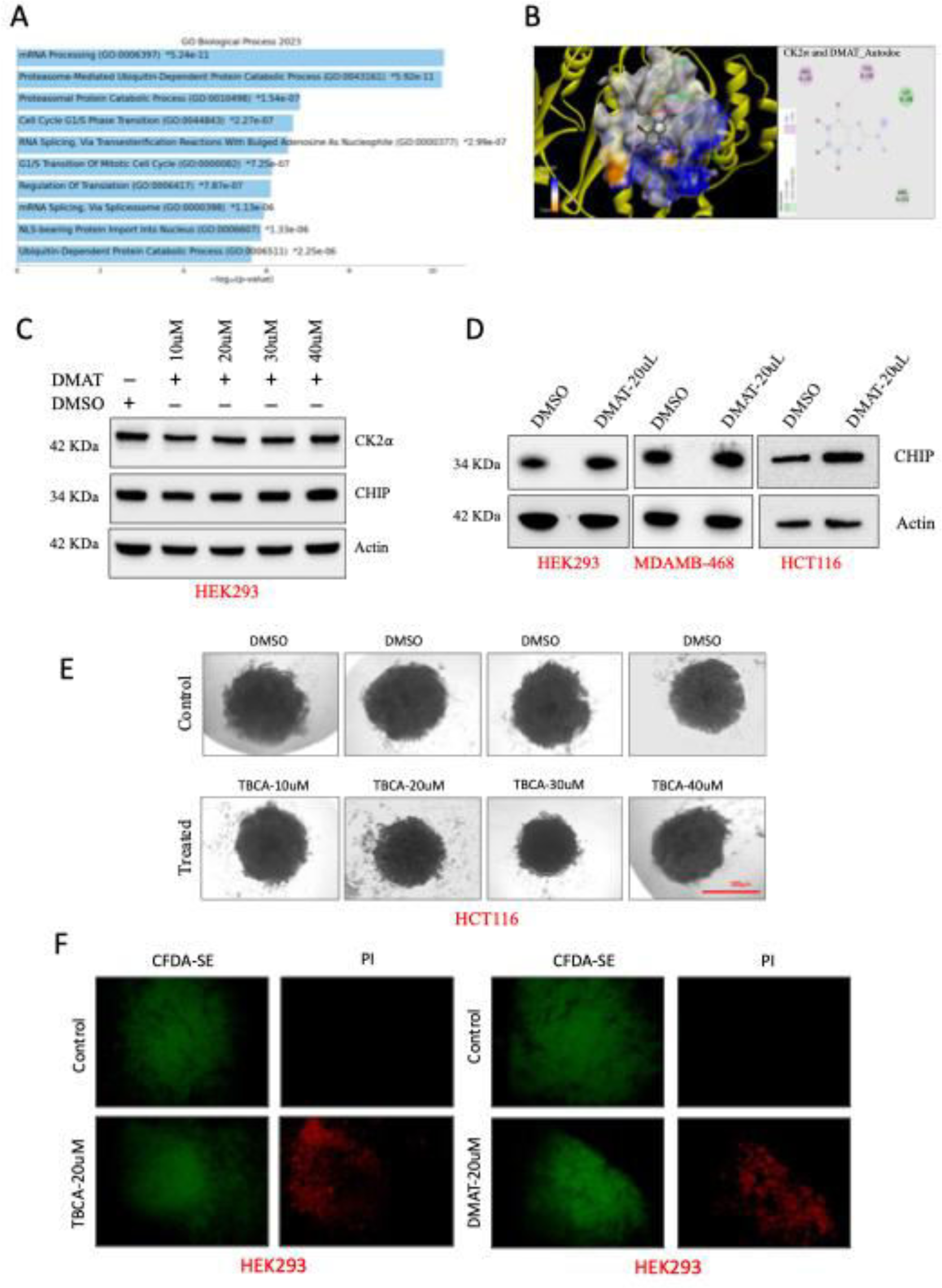
(**A**) Reactome mediated GO: Biological process analysis of CK2α. (**B**) DMAT, a selective inhibitor of CK2α binding sites visualization after docking with Autodoc. (**C**) Immunoblot analysis of HEK293 cells treated with increasing concentrations of DMAT, a selective inhibitor of CK2α (10-40 μM) shows a dose-dependent increase in CHIP protein levels, confirming CK2α inhibition enhances CHIP stability. (**D**) Western blot analysis of HEK293, MDAMB-468, and HCT116 cells treated with DMAT (20 μM) further demonstrates CHIP stabilization compared with DMSO controls. (**E**) Morphological assessment of HCT116 spheroids reveals disrupted architecture and reduced in size upon TBCA treatment compared with DMSO controls, confirming impaired tumorigenic potential. (**F**) Live-dead staining of HEK293 spheroids treated with TBCA (20 μM) shows reduced CFDA-SE (live cell marker, green) and increased PI (dead cell marker, red) signals, indicating apoptosis induction.

**Supplementary Fig. 4.**
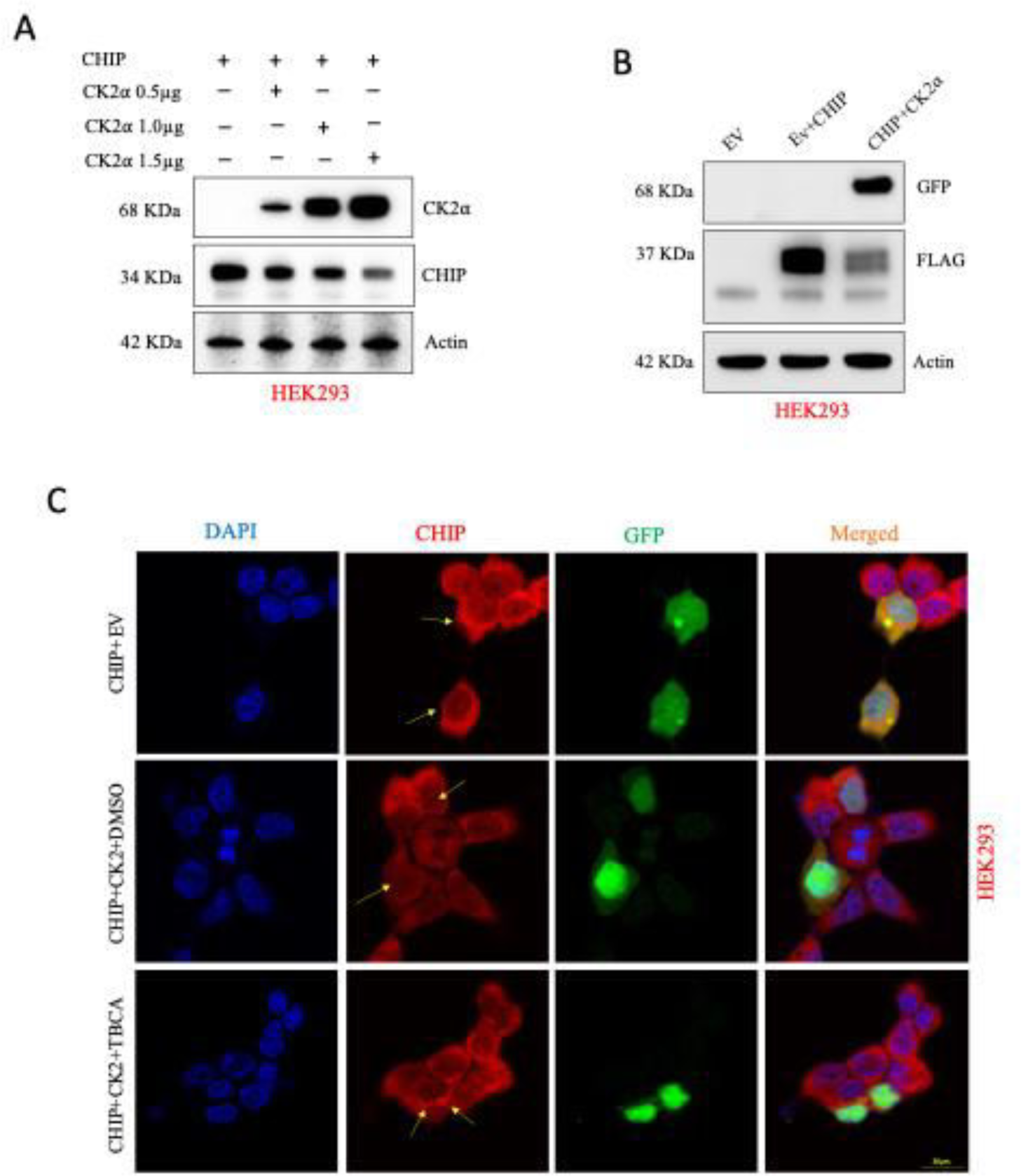
(**A**) Immunoblot analysis of HEK293 cells transfected with empty vector (EV), CHIP, or CHIP + CK2α confirms reduced CHIP protein levels in the presence of CK2α, consistent with CK2α-mediated CHIP destabilization. (**B**) Dose-dependent immunoblot analysis of HEK293 cells transfected with increasing concentrations of CK2α (0.5-1.5 μg) demonstrates progressive CHIP reduction, validating CK2α as a negative regulator of CHIP stability. (**C**) Immunofluorescence imaging of HEK293 cells co-transfected with CHIP and CK2α shows reduced CHIP intensity under DMSO control, whereas TBCA treatment restores CHIP expression and nuclear localization (DAPI, blue; CHIP, red; GFP-tagged CK2α, green; scale bar: 20 μM).

**Supplementary Fig. 5.**
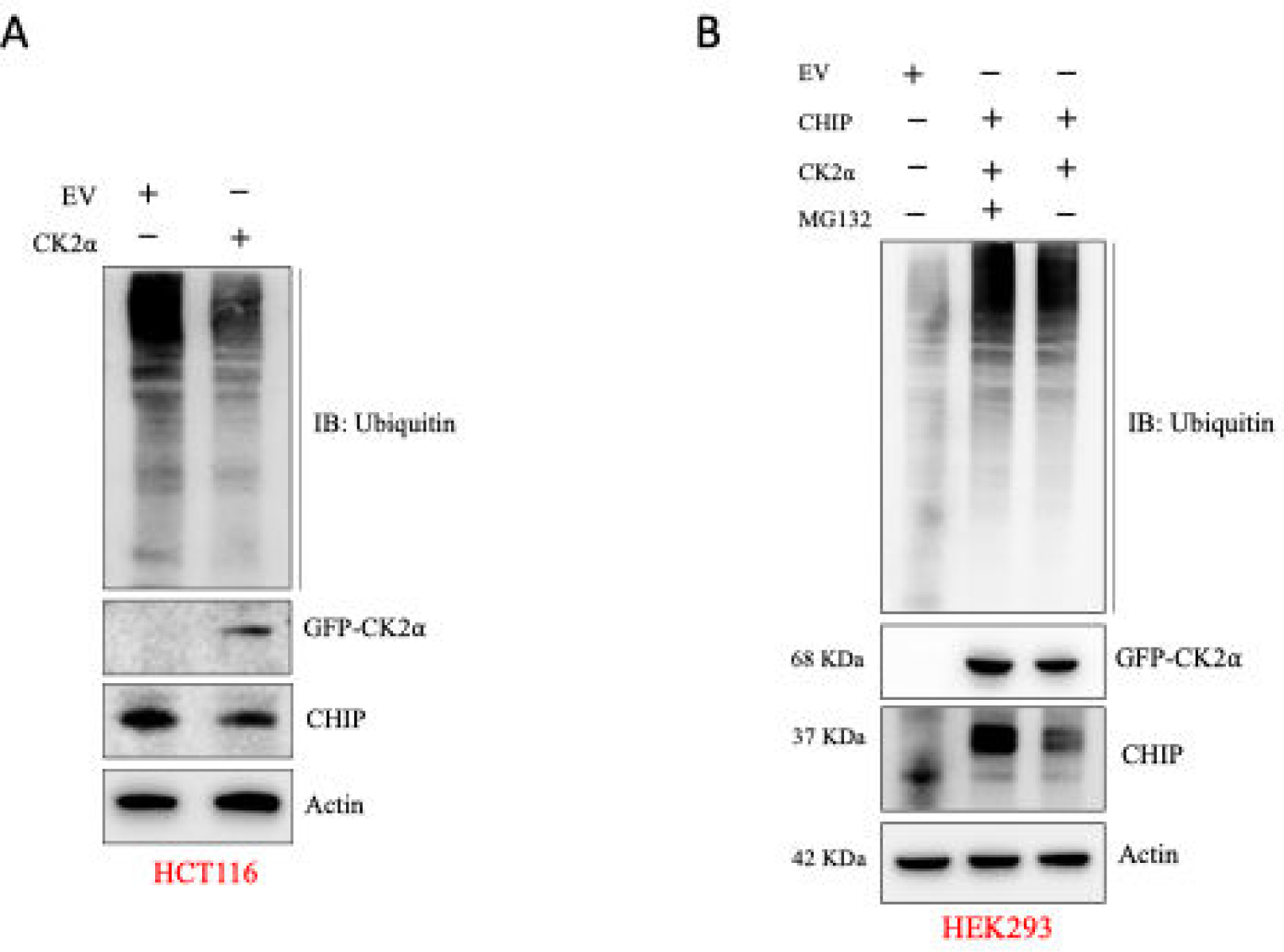
(**A**) HCT116 cells were transfected with GFP-tagged CK2α, with or without CHIP, and cell lysates were immunoblotted with anti-ubiquitin antibody. Actin was used as a loading control. (**B**) HEK293 cells were transfected with CK2α in the presence or absence of CHIP, and ubiquitination levels were examined by immunoblotting. Treatment with the proteasome inhibitor MG132 was included to assess ubiquitinated CK2α accumulation. Molecular weights are indicated to the left of each blot.

**Supplementary Fig. 6.**
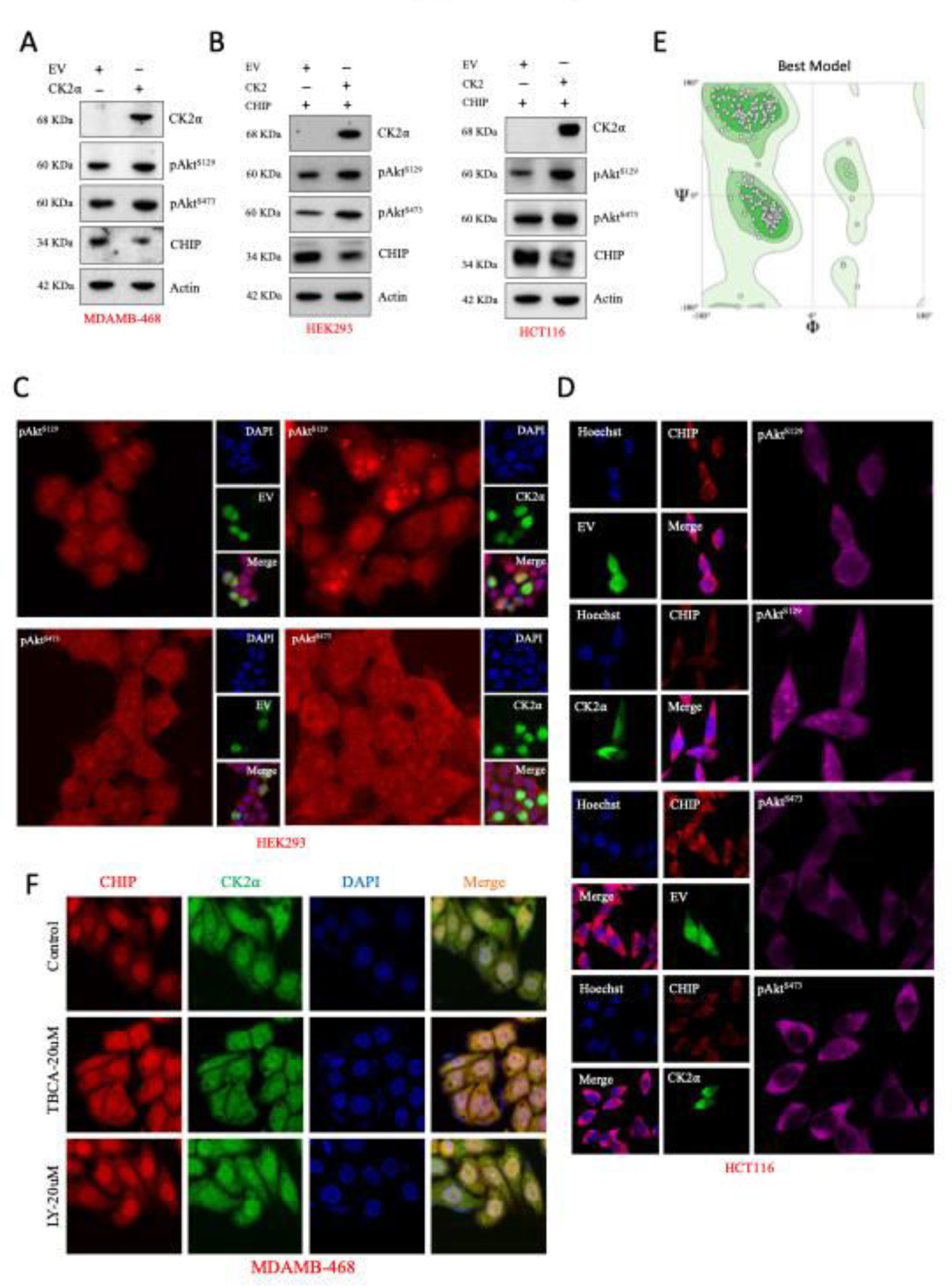
(**A-B**) Immunoblot analysis of Akt phosphorylation in MDA-MB-468 (A), HEK293 and HCT116 (B), cells. Cells were transfected with CK2α in the presence or absence of CHIP. Levels of Akt phosphorylation at Ser129 (pAkt^S129^) and Ser473 (pAkt^S473^) were detected, with actin used as a loading control. (**C-D**) Immunofluorescence analysis of Akt phosphorylation in HEK293 and HCT116 cells. Cells were transfected with CK2α and/or CHIP, and stained for pAkt^S129^or pAkt^S473^ (red), CK2α or CHIP (green), and nuclei (Hoechst/DAPI, blue). Merged images show co-localization. (**E**) Ramachandran plot showing favourable amino acids of Akt and CHIP docked structure of ClusPro best model. (**F**) In MDAMB-468 pharmacological inhibition of CK2α activity with LY-20 μM or TBCA-20 μM increases CHIP level.

**Supplementary Fig. 7.**
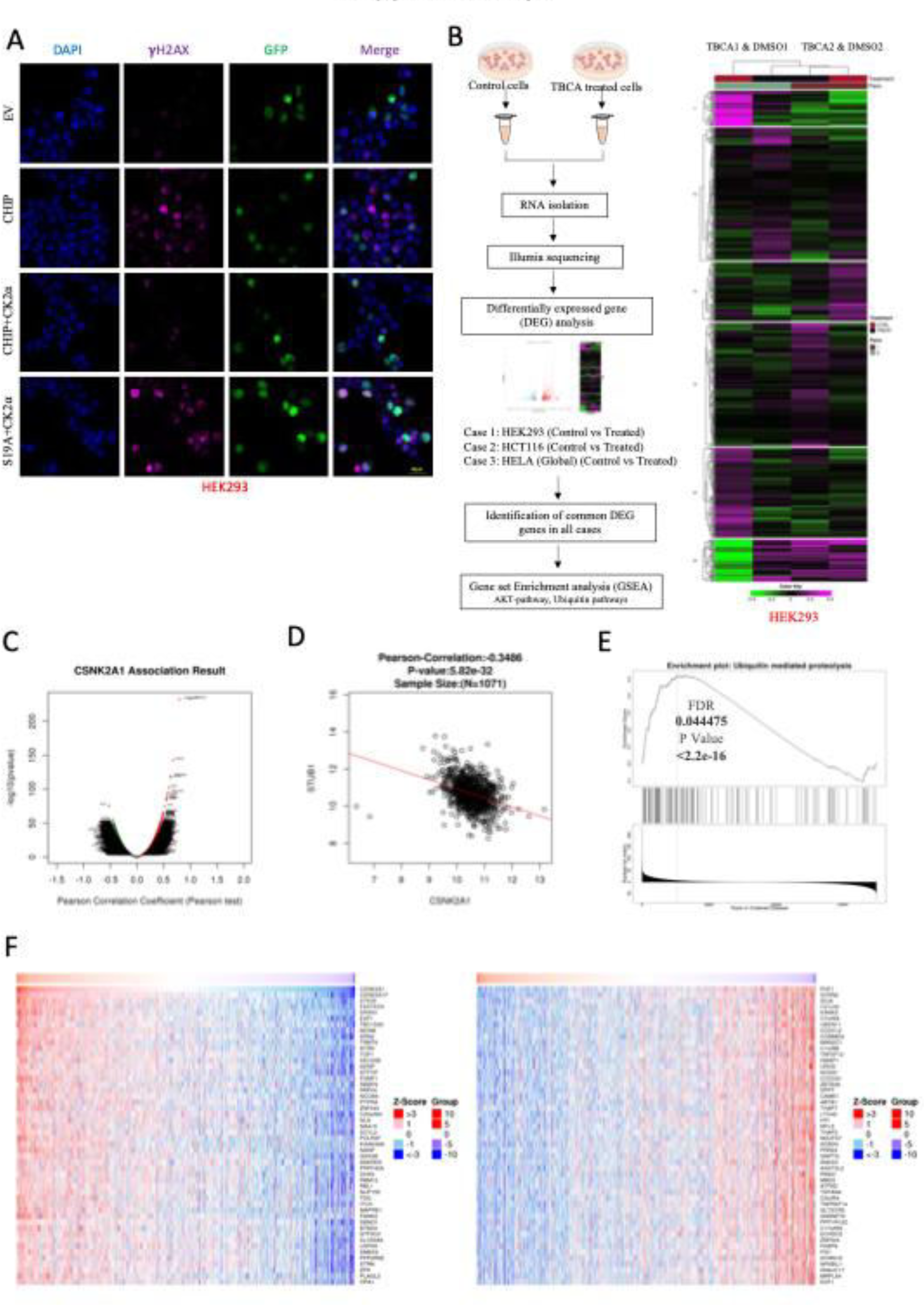
(**A**) Validation of the CK2α-CHIP axis in HEK293 cells by immunofluorescence. Cells were transfected with EV, CHIP, CK2α, or CHIP-S19A mutant construct, stained for γH2AX (DNA damage marker, red), GFP/CK2α or CHIP (green), and nuclei (DAPI, blue). Representative merged images demonstrate altered DNA damage response under conditions of CK2α overexpression, however this function was altered in CHIP (S19A). (**B**) Schematic overview of the RNA-sequencing workflow. Control and TBCA-treated cells (HEK293, HCT116, and one globally available HeLa cell data) were subjected to RNA isolation, Illumina sequencing, and differentially expressed gene (DEG) analysis. Common DEGs across the three cell types were identified, followed by Gene Set Enrichment Analysis (GSEA), which revealed enrichment of Akt-and ubiquitin-related pathways. (**C**) LinkedOmics analysis shows *CSNK2A1* (CK2α) expression association in TCGA-BRCA cohort, followed by DEG analysis. (**D**) Negative Pearson-correlation (-0.34) between *CSNK2A1* and *STUB1* (CHIP) expression confirmed in TCGA-BRCA dataset. (**E**) Also our GSEA analysis showed statistically significant gene expression changes in ubiquitin mediated pathways with FDR = 0.044, P < 2.2e−16. (**F**) Top 50 upregulated and downregulated genes correlated with CSNK2A1 expression in breast cancer.

## Supplementary Tables

**Table S1:** List of antibodies used in this study.

| Antibody | Dilution | Catalogue # | Source |
| --- | --- | --- | --- |
| CK2α | 1:2000 (WB) 1:500 (ICC/IF) | 2656S | Cell Signaling Technology |
| CK2α | 1:2000 (WB)<br>1:50 (ICC/IF) | sc-12738 | Santacruz Biotechnology |
| CHIP | 1:50 (ICC/IF) | ab84781 | AbCam |
| CHIP | 1:1000 (WB) | sc-133066 | Santacruz Biotechnology |
| p-CHIP (Ser19) | 1:1000 (WB)<br>1:50 (IF) | AF3514 | Affinity Biosciences |
| Actin | 1:2000 (WB) | A5441 | Sigma |
| GFP | 1:2000 | G6539 | Sigma |
| GFP | 1:2000 | sc-9996 | Santacruz Biotechnology |
| FLAG | 1:2000 (WB) 1:100 (IP) | F7424 | Sigma |
| Bcl-xL | 1:2000 | 2764 | Cell Signaling Technology |
| Akt | 1:1000 (WB) | 9272 | Cell Signalling Technology |
| pAkt (S473) | 1:1000 (WB) | 9271 | Cell Signalling Technology |
| pAkt (S129) | 1:500 (WB)<br>1:500 (ICC) | PA5-12540 | Thermo Scientific |
| Bax | 1:1000 (WB) | 2774 | Cell Signalling Technology |
| Bim | 1:1000 (WB) | 2819 | Cell Signalling Technology |
| Ubiquitin (P4D1) | 1:1000 (WB)<br>1:100 (IP) | sc-8017 | Santa Cruz Biotechnology |
| PCNA (PC10) | 1:1000 (WB)<br>1:50 (ICC) | 2586 | Cell Signalling Technology |
| Caspase-3 | 1:1000 (WB) | 9662 | Cell Signaling Technology |
| Cleaved Caspase-3 | 1:1000 (WB) | 9664 | Cell Signaling Technology |
| Alexafluor647- tagged anti-γH2AX | 1:50 (ICC) | 560447 | BD Pharmingen |
| Anti-Rabbit IgG AF488 | 1:500 (ICC/IF) | A11008 | Molecular Probes |
| Anti-Rabbit IgG AF594 | 1:500 (ICC/IF) | A11012 | Molecular Probes |
| Anti-Mouse IgG AF488 | 1:500 (ICC/IF) | A11001 | Molecular Probes |
| Anti-Mouse IgG AF594 | 1:500 (ICC/IF) | A11005 | Molecular Probes |
| Anti-Goat IgG AF488 | 1:500 (ICC) | A11055 | Molecular Probes |
| Anti-Goat IgG AF594 | 1:500 (ICC) | A11058 | Molecular Probes |
| Normal Rabbit IgG | 1:500 (IP) | sc-2027 | Santa Cruz Biotechnology |
| Normal Mouse IgG | 1:500 (IP) | sc-2025 | Santa Cruz Biotechnology |
| Normal Goat IgG | 1:500 (IP) | sc-2028 | Santa Cruz Biotechnology |
| Anti - goat secondary | 1:1000 (WB) | A5420 | Sigma |
| Anti - rabbit secondary | 1:1000 (WB) | A0545 | Sigma |
| Anti - mouse secondary | 1:1000 (WB) | A2554 | Sigma |

**Table S2:** List of online tools and software used

| Name | Link |
| --- | --- |
| cBioPortal | https://www.cbioportal.org/ |
| Image Lab | https://www.bio-rad.com/en-in/product/image-lab-software?ID=KRE6P5E8Z |
| Chimera | https://www.cgl.ucsf.edu/chimera/download.html |
| ClusPro 2.0 | https://cluspro.org/help.php |
| Cytoscape | http://cytoscape.org |
| Ensembl | https://www.ensembl.org/Homo_sapiens/Info/Index |
| GEPIA | http://gepia.cancer-pku.cn/ |
| GeneMANIA | http://genemania.org |
| GeneMiner | https://github.com/happywithxpl/GeneMiner |
| GraphPad Prism | https://www.graphpad.com |
| Galaxy | https://usegalaxy.org/ |
| GPS 6.0 | https://gps.biocuckoo.cn/online.php |
| Inkscape | https://inkscape.org/ |
| Jalview | https://www.jalview.org/ |
| KEGG Database | https://www.genome.jp/kegg/ |
| MaxQuant | https://www.maxquant.org/ |
| MEGA | https://www.megasoftware.net/ |
| Mendeley | https://www.mendeley.com/ |
| NCBI Primer Blast | https://www.ncbi.nlm.nih.gov/tools/primer-blast/ |
| PyMOL | https://www.pymol.org/ |
| QuikChange Primer Design | https://www.agilent.com/store/primerDesignProgram.jsp |
| RCSB PDB | https://www.rcsb.org/ |
| SnapGene | https://www.snapgene.com/ |
| Single Cell Expression Atlas | https://www.ebi.ac.uk/gxa/sc/home |
| TCGA Dataset | https://www.cancer.gov/ccg/research/genome-sequencing/tcga |

**Table S3:**
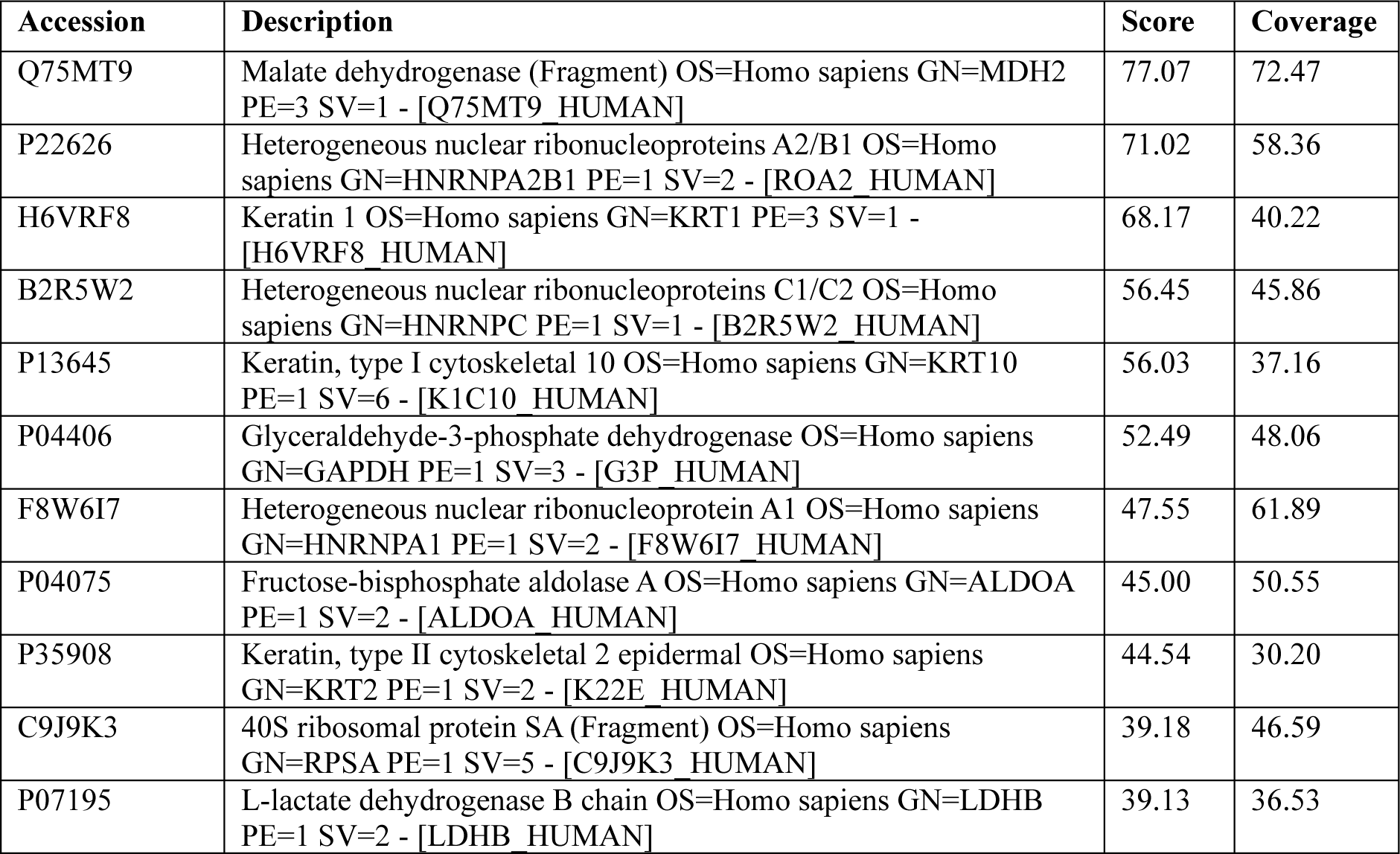
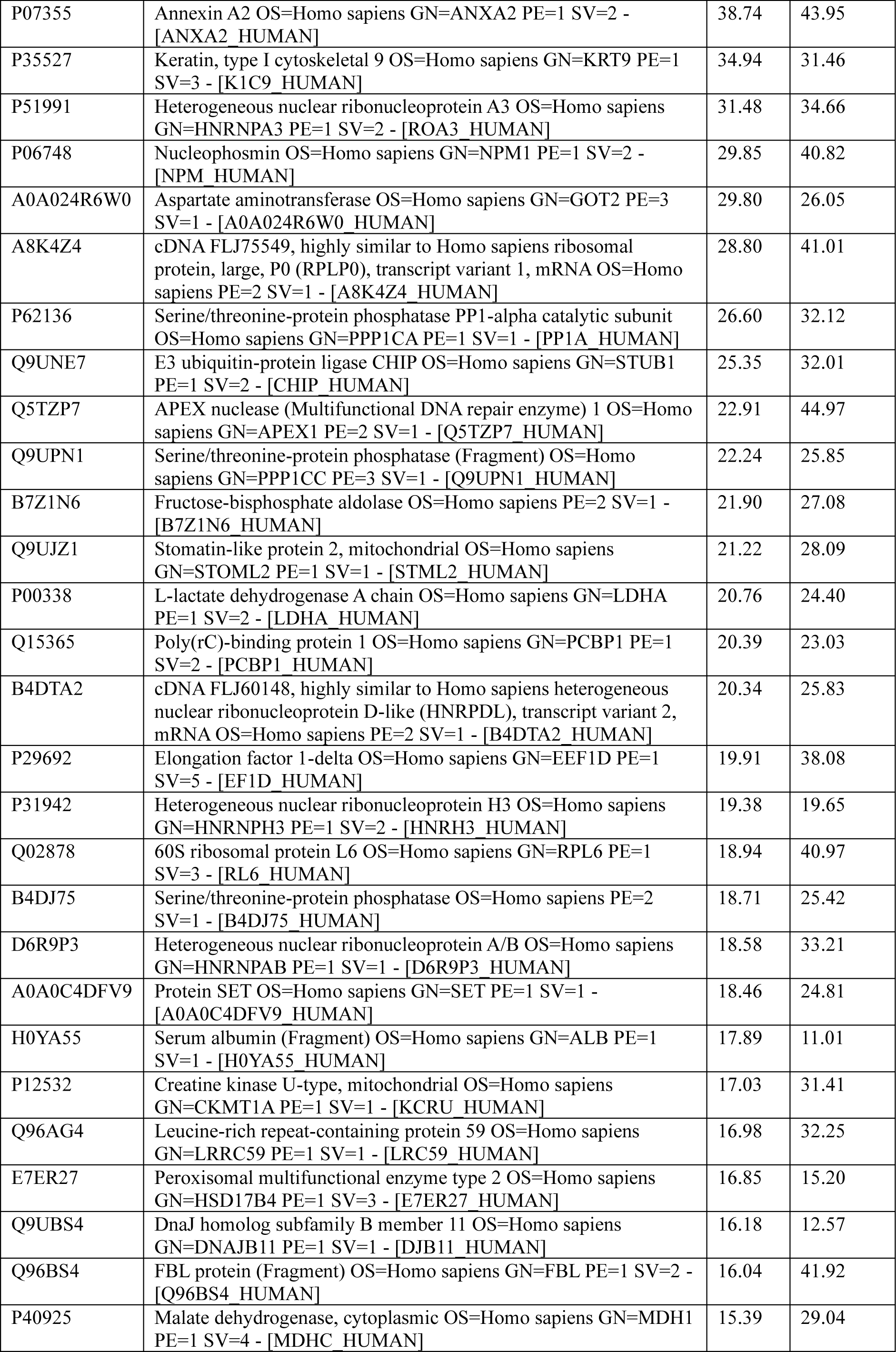
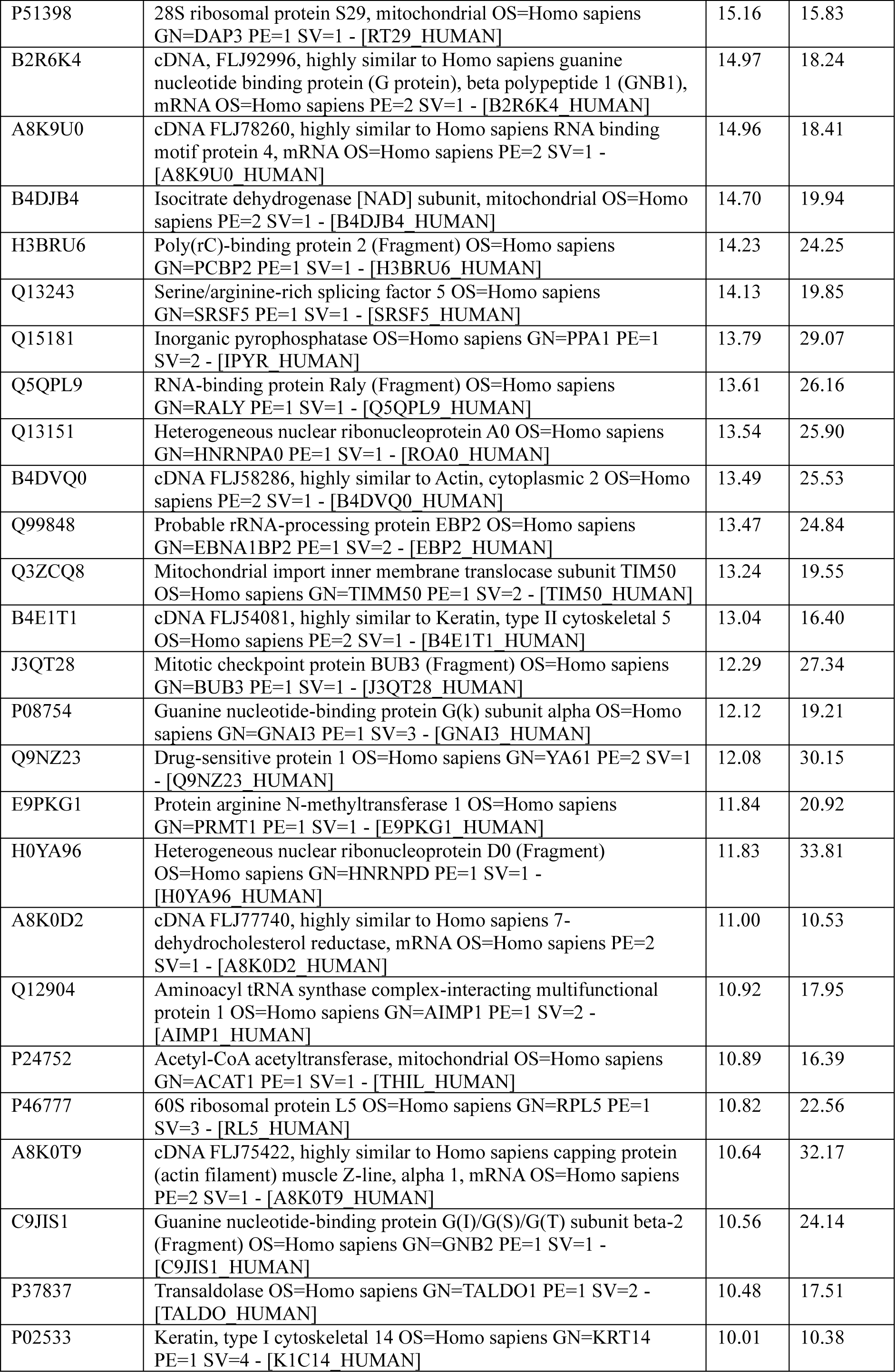
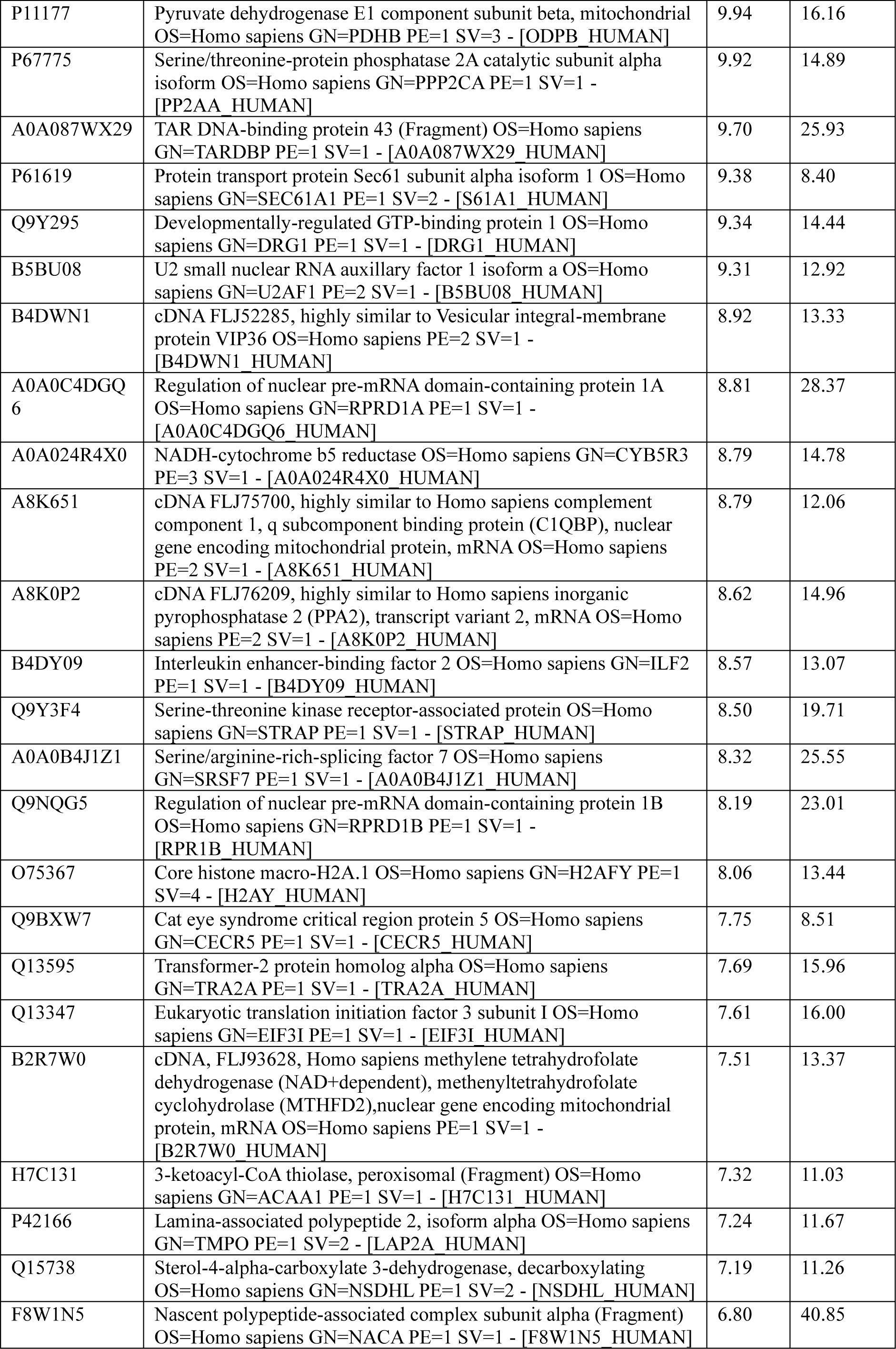
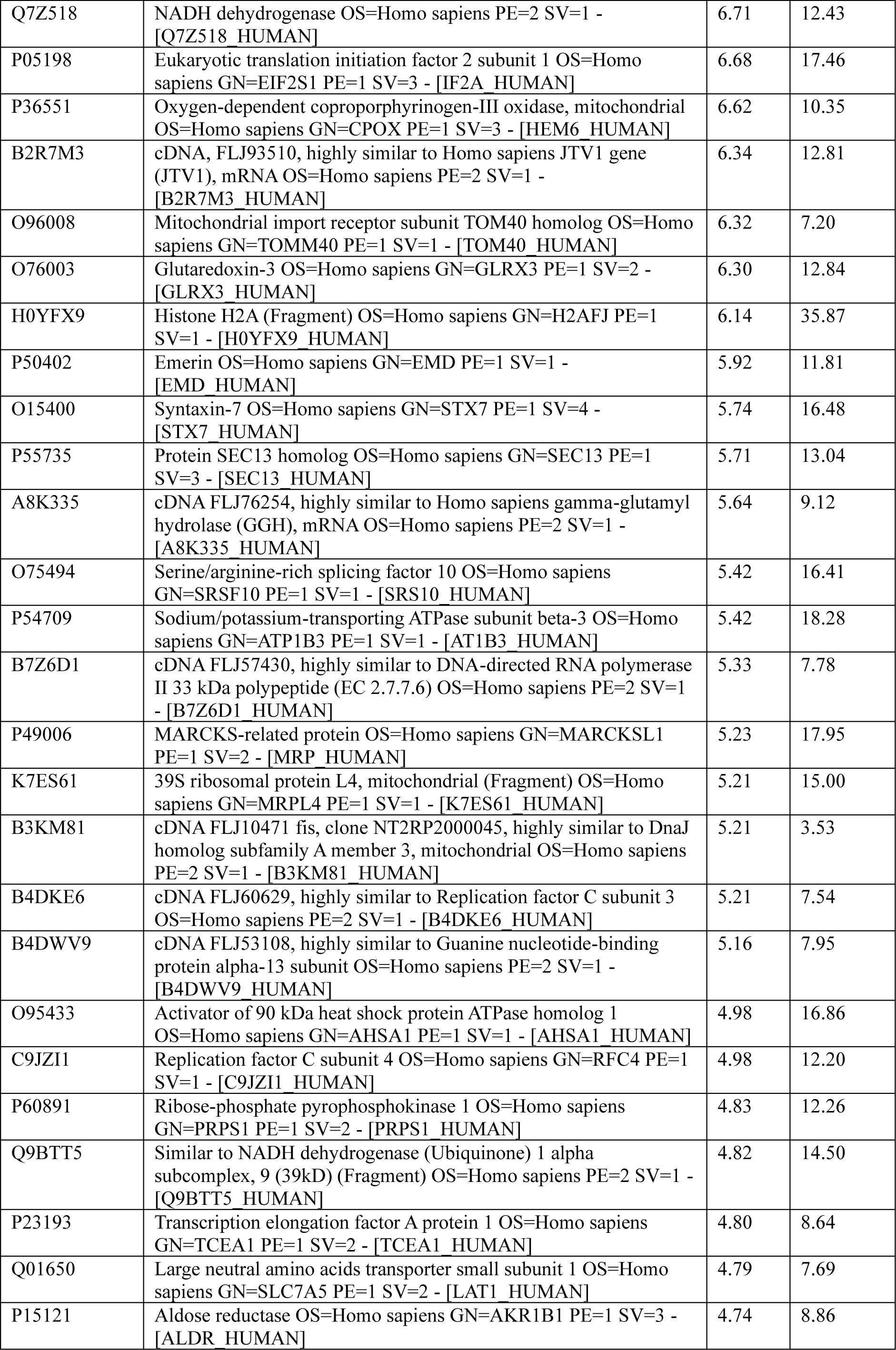
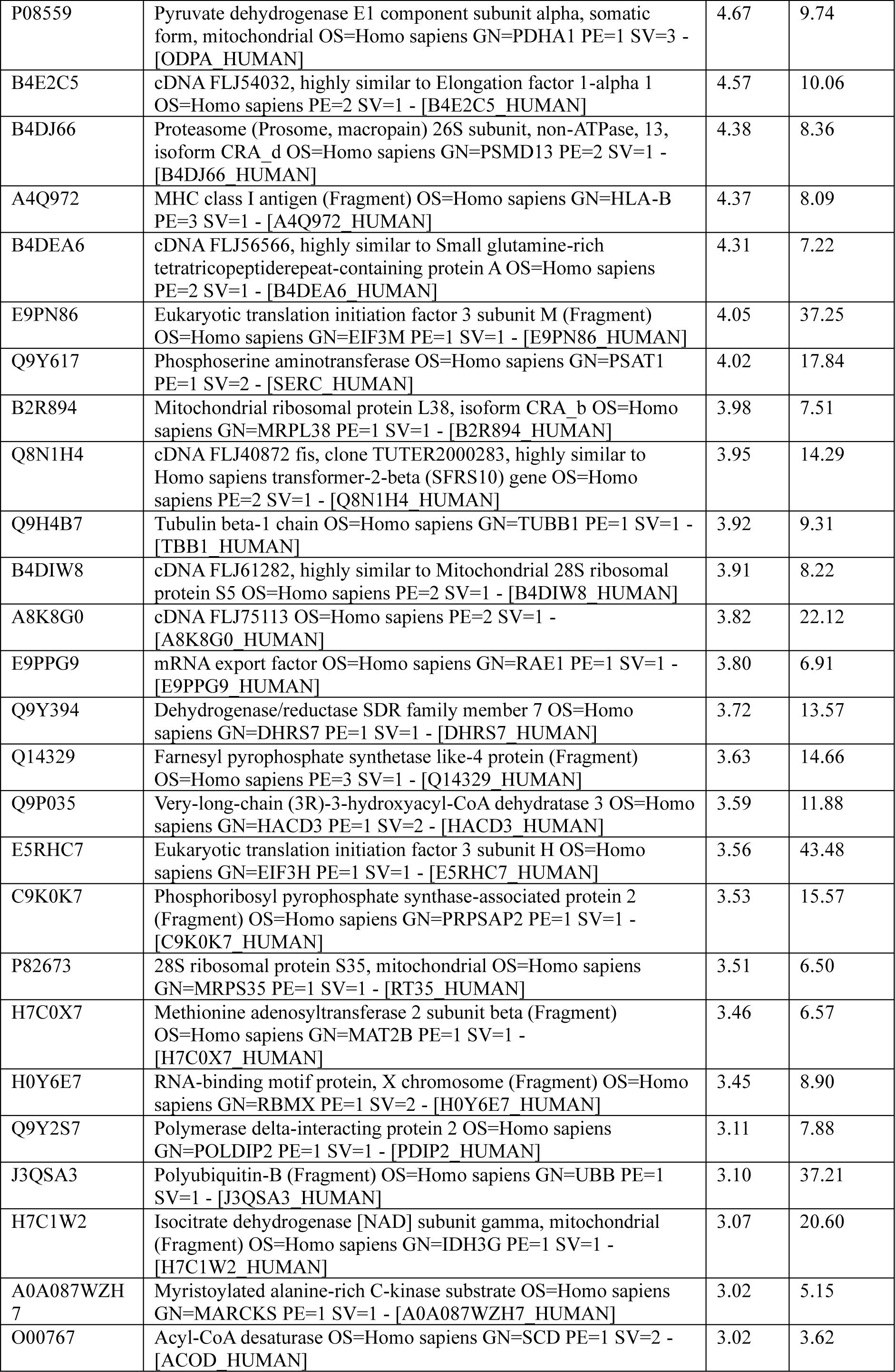
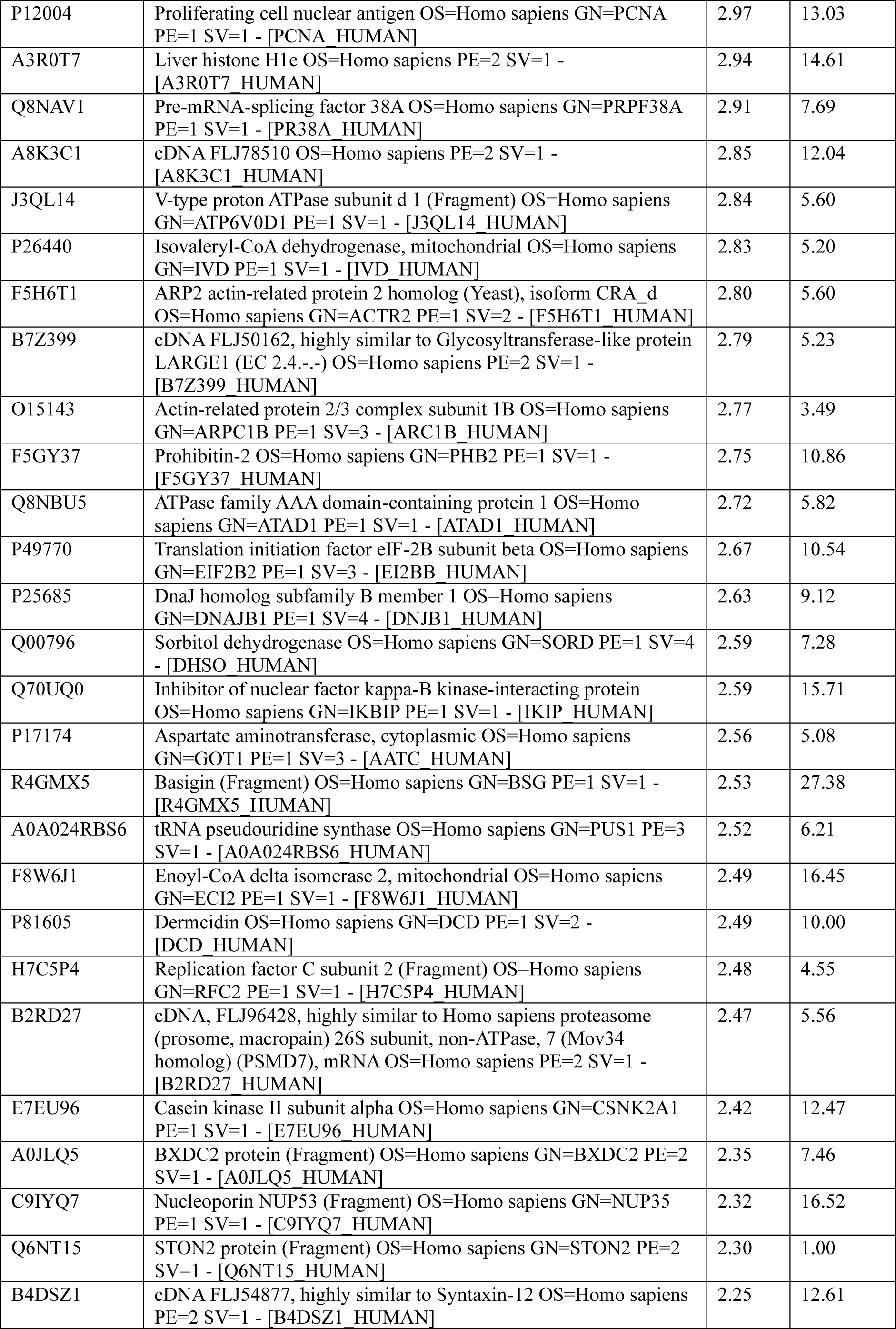
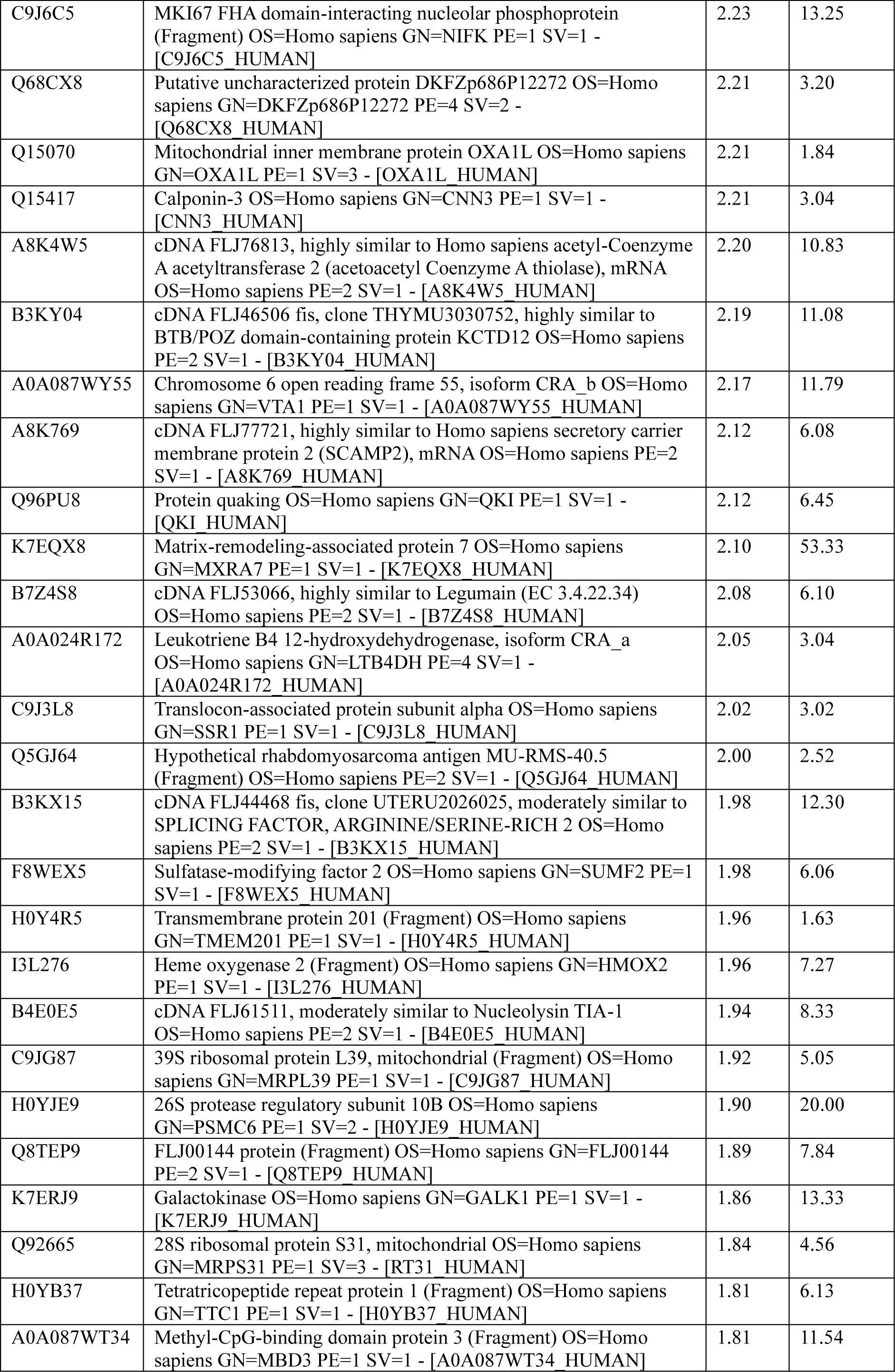
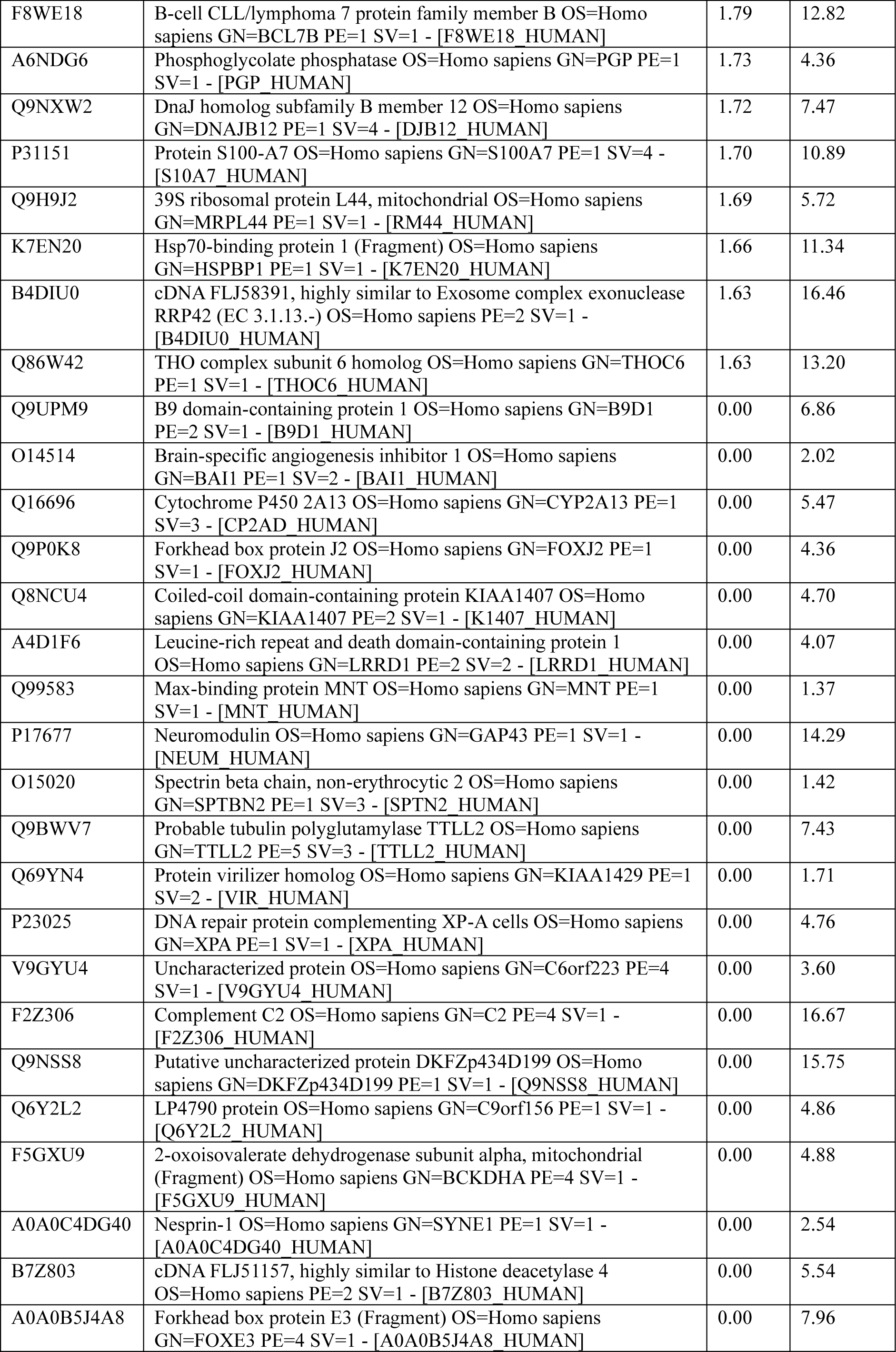
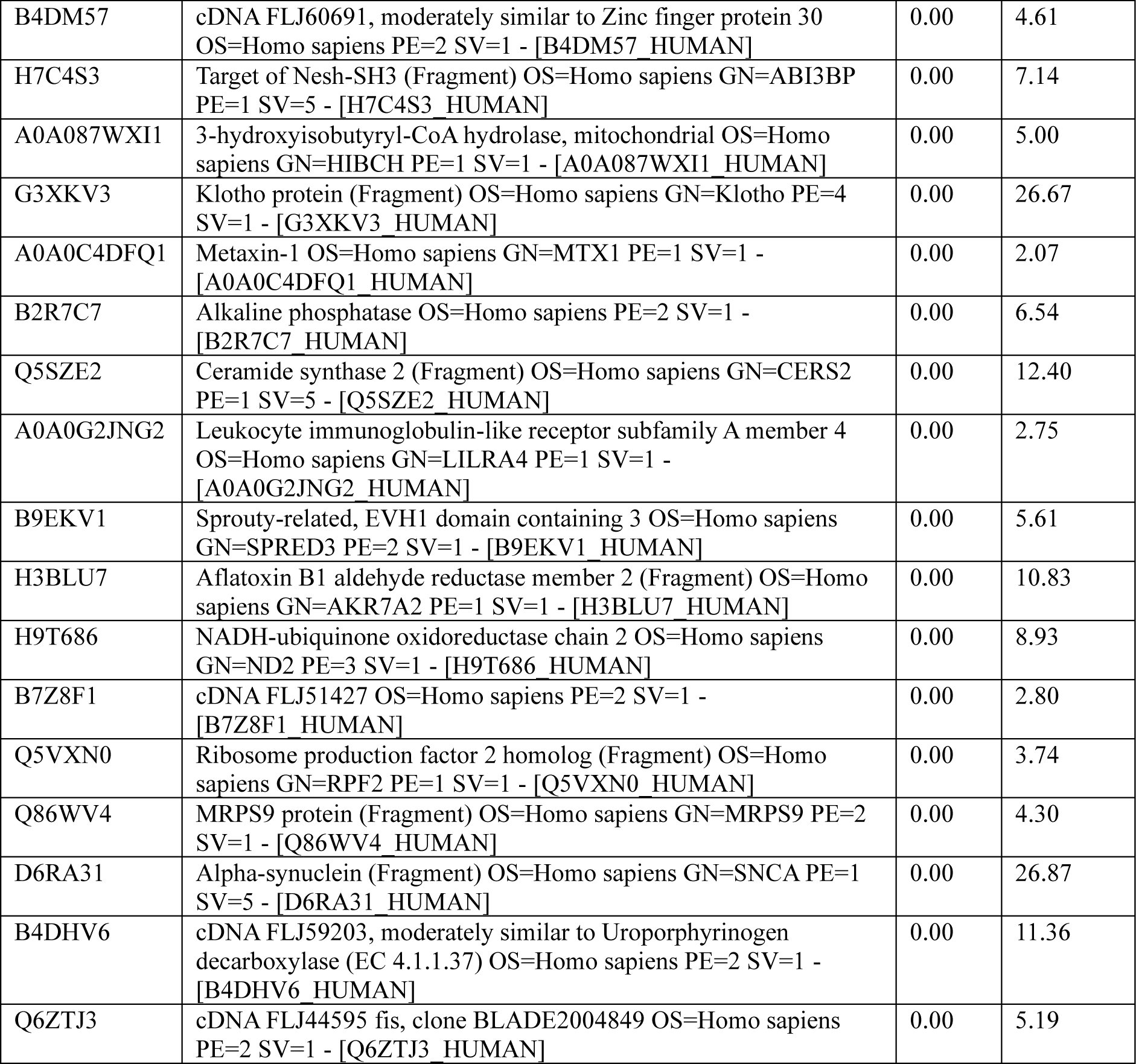
List of Protein identified after LC-MS

**Table S4:**
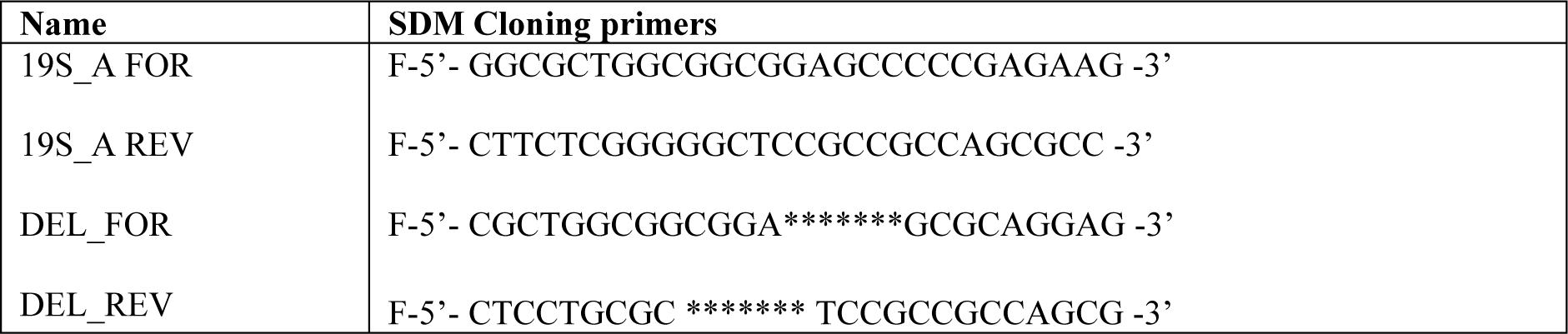
List of primers used

## REFERENCES

1. Murata S, Minami Y, Minami M, Chiba T, Tanaka K. CHIP is a chaperone-dependent E3 ligase that ubiquitylates unfolded protein. EMBO Rep 2001; 2. doi:10.1093/embo-reports/kve246.

2. Wang Y, Ren F, Wang Y, Feng Y, Wang D, Jia B et al. CHIP/Stub1 functions as a tumor suppressor and represses NF-κB-mediated signaling in colorectal cancer. Carcinogenesis 2014; 35. doi:10.1093/carcin/bgt393.

3. Wang T, Yang J, Xu J, Li J, Cao Z, Zhou L et al. CHIP is a novel tumor suppressor in pancreatic cancer through targeting EGFR. Oncotarget 2014; 5. doi:10.18632/oncotarget.1890.

4. Jin B, Wang M, Sun Y, Lee PAH, Zhang X, Lu Y et al. CHIP suppresses the proliferation and migration of A549 cells by mediating the ubiquitination of eIF2α and upregulation of tumor suppressor RBM5. Journal of Biological Chemistry 2024; 300. doi:10.1016/j.jbc.2024.105673.

5. Wang S, Wu X, Zhang J, Chen Y, Xu J, Xia X et al. CHIP functions as a novel suppressor of tumour angiogenesis with prognostic significance in human gastric cancer. Gut 2013; 62. doi:10.1136/gutjnl-2011-301522.

6. Ko HR, Kim CK, Lee SB, Song J, Lee KH, Kim KK et al. P42 Ebp1 regulates the proteasomal degradation of the p85 regulatory subunit of PI3K by recruiting a chaperone-E3 ligase complex HSP70/CHIP. Cell Death Dis 2014; 5. doi:10.1038/cddis.2014.79.

7. Paul I, Ahmed SF, Bhowmik A, Deb S, Ghosh MK. The ubiquitin ligase CHIP regulates c-Myc stability and transcriptional activity. Oncogene 2013; 32. doi:10.1038/onc.2012.144.

8. Paul I, Ghosh MK. A CHIPotle in physiology and disease. International Journal of Biochemistry and Cell Biology. 2015; 58. doi:10.1016/j.biocel.2014.10.027.

9. Trembley JH, Wang G, Unger G, Slaton J, Ahmed K. Protein kinase CK2 in health and disease: CK2: a key player in cancer biology. Cell Mol Life Sci 2009; 66.

10. Guerra B, Issinger OG. Protein kinase CK2 and its role in cellular proliferation, development and pathology. Electrophoresis. 1999; 20. doi:10.1002/(SICI)1522-2683(19990201)20:2<391::AID-ELPS391>3.0.CO;2-N.

11. Niefind K, Issinger OG. Primary and secondary interactions between CK2α and CK2β lead to ring-like structures in the crystals of the CK2 holoenzyme. Mol Cell Biochem 2005; 274. doi:10.1007/s11010-005-3114-0.

12. Ahmad KA, Wang G, Unger G, Slaton J, Ahmed K. Protein kinase CK2 - A key suppressor of apoptosis. Adv Enzyme Regul 2008; 48. doi:10.1016/j.advenzreg.2008.04.002.

13. Niefind K, Guerra B, Pinna LA, Issinger OG, Schomburg D. Crystal structure of the catalytic subunit of protein kinase CK2 from Zea mays at 2.1 Å resolution. EMBO Journal 1998; 17. doi:10.1093/emboj/17.9.2451.

14. Karmakar S, Chatterjee M, Basu M, Ghosh MK. CK2: The master regulator in tumor immune-microenvironment - A crucial target in oncotherapy. Eur J Pharmacol 2025; 994: 177376.

15. Dickey CA, Koren J, Zhang YJ, Xu YF, Jinwal UK, Birnbaum MJ et al. Akt and CHIP coregulate tau degradation through coordinated interactions. Proc Natl Acad Sci U S A 2008; 105. doi:10.1073/pnas.0709180105.

16. Su CH, Wang CY, Lan KH, Li CP, Chao Y, Lin HC et al. Akt phosphorylation at Thr308 and Ser473 is required for CHIP-mediated ubiquitination of the kinase. Cell Signal 2011; 23. doi:10.1016/j.cellsig.2011.06.018.

17. Liu Z, Ma L, Wen ZS, Hu Z, Wu FQ, Li W et al. Cancerous inhibitor of PP2A is targeted by natural compound celastrol for degradation in non-small-cell lung cancer. Carcinogenesis 2014; 35. doi:10.1093/carcin/bgt395.

18. Kim C, Yun N, Lee J, Youdim MBH, Ju C, Kim WK et al. Phosphorylation of CHIP at Ser20 by Cdk5 promotes tAIF-mediated neuronal death. Cell Death Differ 2016; 23. doi:10.1038/cdd.2015.103.

19. Ranek MJ, Oeing C, Sanchez-Hodge R, Kokkonen-Simon KM, Dillard D, Aslam MI et al. CHIP phosphorylation by protein kinase G enhances protein quality control and attenuates cardiac ischemic injury. Nat Commun 2020; 11. doi:10.1038/s41467-020-18980-x.

20. Wang D, Qian X, Nancy Du Y-C, Sanchez-Solana B. cProSite: A Web Based Interactive Platform for Online Proteomics, Phosphoproteomics, and Genomics Data Analysis. Journal of Biotechnology and Biomedicine 2023; 06. doi:10.26502/jbb.2642-91280119.

21. Chandrashekar DS, Karthikeyan SK, Korla PK, Patel H, Shovon AR, Athar M et al. UALCAN: An update to the integrated cancer data analysis platform. Neoplasia (United States) 2022; 25. doi:10.1016/j.neo.2022.01.001.

22. Tang G, Cho M, Wang X. OncoDB: An interactive online database for analysis of gene expression and viral infection in cancer. Nucleic Acids Res 2022; 50. doi:10.1093/nar/gkab970.

23. Pagano MA, Poletto G, Di Maira G, Cozza G, Ruzzene M, Sarno S et al. Tetrabromocinnamic acid (TBCA) and related compounds represent a new class of specific protein kinase CK2 inhibitors. ChemBioChem 2007; 8. doi:10.1002/cbic.200600293.

24. Ryu SY, Kim S. Evaluation of CK2 inhibitor (E)-3-(2,3,4,5-tetrabromophenyl)acrylic acid (TBCA) in regulation of platelet function. Eur J Pharmacol 2013; 720. doi:10.1016/j.ejphar.2013.09.064.

25. Chatterjee A, Chatterjee U, Ghosh MK. Activation of protein kinase CK2 attenuates FOXO3a functioning in a PML-dependent manner: implications in human prostate cancer. Cell Death Dis 2013; 4: e543–e543.

26. Schneider CC, Hessenauer A, Götz C, Montenarh M. DMAT, an inhibitor of protein kinase CK2 induces reactive oxygen species and DNA double strand breaks. Oncol Rep 2009; 21. doi:10.3892/or_00000392.

27. Pagano MA, Meggio F, Ruzzene M, Andrzejewska M, Kazimierczuk Z, Pinna LA. 2-Dimethylamino-4,5,6,7-tetrabromo-1H-benzimidazole: A novel powerful and selective inhibitor of protein kinase CK2. Biochem Biophys Res Commun 2004; 321. doi:10.1016/j.bbrc.2004.07.067.

28. Paul I, Ghosh MK. The E3 ligase CHIP: Insights into its structure and regulation. Biomed Res Int. 2014; 2014. doi:10.1155/2014/918183.

29. Das N, Datta N, Chatterjee U, Ghosh MK. Estrogen receptor alpha transcriptionally activates casein kinase 2 alpha: A pivotal regulator of promyelocytic leukaemia protein (PML) and AKT in oncogenesis. Cell Signal 2016; 28. doi:10.1016/j.cellsig.2016.03.007.

30. Cao Z, Li G, Shao Q, Yang G, Zheng L, Zhang T et al. CHIP: A new modulator of human malignant disorders. Oncotarget 2016; 7. doi:10.18632/oncotarget.8219.

31. Di Maira G, Salvi M, Arrigoni G, Marin O, Sarno S, Brustolon F et al. Protein kinase CK2 phosphorylates and upregulates Akt/PKB. Cell Death Differ 2005; 12. doi:10.1038/sj.cdd.4401604.

32. Saha G, Sarkar S, Mohanta PS, Kumar K, Chakrabarti S, Basu M et al. USP7 targets XIAP for cancer progression: Establishment of a p53-independent therapeutic avenue for glioma. Oncogene 2022; 41. doi:10.1038/s41388-022-02486-5.

33. Chakraborty S, Karmakar S, Basu M, Kal S, Ghosh MK. The E3 ubiquitin ligase CHIP drives monoubiquitylation-mediated nuclear import of the tumor suppressor PTEN. J Cell Sci 2023; 136. doi:10.1242/jcs.260950.

34. Shaw R, Karmakar S, Basu M, Ghosh MK. DDX5 (p68) orchestrates β-catenin, RelA and SP1 mediated MGMT gene expression in human colon cancer cells: Implication in TMZ chemoresistance. Biochim Biophys Acta Gene Regul Mech 2023; 1866. doi:10.1016/j.bbagrm.2023.194991.

35. Kal S, Chakraborty S, Karmakar S, Ghosh MK. Wnt/β-catenin signaling and p68 conjointly regulate CHIP in colorectal carcinoma. Biochim Biophys Acta Mol Cell Res 2022; 1869. doi:10.1016/j.bbamcr.2021.119185.

36. Sarkar M, Khare V, Guturi KKN, Das N, Ghosh MK. The DEAD box protein p68: A crucial regulator of AKT/FOXO3a signaling axis in oncogenesis. Oncogene 2015; 34. doi:10.1038/onc.2015.42.

37. Białkowska K, Komorowski P, Bryszewska M, Miłowska K. Spheroids as a type of three-dimensional cell cultures—examples of methods of preparation and the most important application. Int J Mol Sci. 2020; 21. doi:10.3390/ijms21176225.

38. Ma HL, Jiang Q, Han S, Wu Y, Tomshine JC, Wang D et al. Multicellular tumor spheroids as an in vivo-like tumor model for three-dimensional imaging of chemotherapeutic and nano material cellular penetration. Mol Imaging 2012; 11. doi:10.2310/7290.2012.00012.

39. Baker BM, Chen CS. Deconstructing the third dimension-how 3D culture microenvironments alter cellular cues. J Cell Sci. 2012; 125. doi:10.1242/jcs.079509.

